# Lewy Body Radius Growth: The Hypothesis of the Cube Root of Time Dependency

**DOI:** 10.1101/2023.10.30.564787

**Authors:** Andrey V. Kuznetsov

## Abstract

This paper presents a model for the growth of Lewy bodies (LBs), which are pathological hallmarks of Parkinson’s disease (PD). The model simulates the growth of classical LBs, consisting of a core and a halo. The core is assumed to comprise lipid membrane fragments and damaged organelles, while the halo consists of radiating alpha-synuclein (α-syn) fibrils. The Finke-Watzky model is employed to simulate the aggregation of lipid fragments and α-syn monomers. By analytically and numerically exploring the solutions of the governing equations, approximate solutions were derived, which are applicable for large times. The application of these approximate solutions to simulate LB radius growth led to the discovery of the cube root hypothesis, which posits that the LB radius is proportional to the cube root of its growth time. Sensitivity analysis revealed that the LB radius is unaffected by the kinetic rates of nucleation and autocatalytic growth, with growth primarily regulated by the production rates of lipid membrane fragments and α-syn monomers. The model suggests that large LBs relevant to PD can only develop when the machinery responsible for degrading lipid membrane fragments, α-syn monomers, and their aggregates is dysfunctional.

## 1. Introduction

Parkinson’s disease (PD) (Cookson, 2005; Armstrong and Okun, 2020; Menšíková et al., 2022), dementia with Lewy bodies (LBs) (Graff-Radford et al., 2017), and other alpha-synucleinopathies (Alafuzoff and Hartikainen, 2017) are neurodegenerative disorders characterized by intracellular inclusions known as LBs. LBs are insoluble, spherical structures that develop within the neuronal soma in these pathological states and have the capacity to spread to various brain regions, and even extend into neural grafts transplanted into the brains of PD patients (Steiner et al., 2011; Dickson and Lin 2014).

Whether LBs are neurotoxic or simply serve as a mechanism to clear damaged organelles remains unclear (Parkkinen et al., 2011; Shahmoradian et al., 2019; Lashuel, 2020; Fares et al., 2021; Power et al., 2017; Olanow et al., 2004). According to the findings reported in Parkkinen et al. (2011), in PD, there is a significant loss of neurons in the ventrolateral tier of the pars compacta of the substantia nigra (SN). This region houses dopaminergic neurons, which are the primary neurons affected in PD. The extent of neuron loss in this region surpasses the number of Lewy bodies (LBs) present. On the other hand, as PD advances, the number of neurons containing LBs in the SN pars compacta does not increase; instead, LB-laden neurons undergo cell death (Shults, 2006).

Classical LBs consist of a dense core surrounded by a halo of radiating filaments (Shahmoradian et al., 2019; Lashuel, 2020). For a long time LBs were believed to primarily consist of alpha-synuclein (α-syn) fibrils. This view was well supported by numerous experimental studies (Spillantini et al., 1998; Mahul-Mellier et al., 2020; Lashuel 2020). Recently Shahmoradian et al. (2019) reported that LBs are composed of lipid membrane fragments, altered organelles (such as damaged or distorted lysosomes and mitochondria), and non-fibrillar forms of α-syn. This surprising observation has sparked intense debates regarding the inclusion of α-syn fibrils within LBs (Lashuel, 2020; Fares et al. 2021).

To address these contradictory findings, the model developed here posits a two-stage process in the formation of LBs. Classical brainstem-type LBs consist of a dense core at their center and a surrounding halo composed of radiating filaments (Mahul-Mellier et al., 2020; Fares et al., 2021). Both the core and the halo have spherical shapes. It is probable that LBs originate from pale bodies, which are granular structures without a halo (Fares et al., 2021). A pale body can transform into a classical LB with a halo (Shults, 2006).

The model proposes a two-stage process. In the first stage, the core of an LB is formed through aggregation of lipid membrane fragments and altered organelles. Subsequently, surface-initiated polymerization (Gambinossi et al., 2015), prompted by the core’s surface, triggers the growth of α-syn fibrils, which form the LB’s halo (Fig. 1). The α-syn fibrils extend by incorporating α-syn monomers from the cytosol. These radiating filaments also act as a barrier, inhibiting further core growth by preventing sequestration of lipid-rich material by the core.

**Fig. 1.**
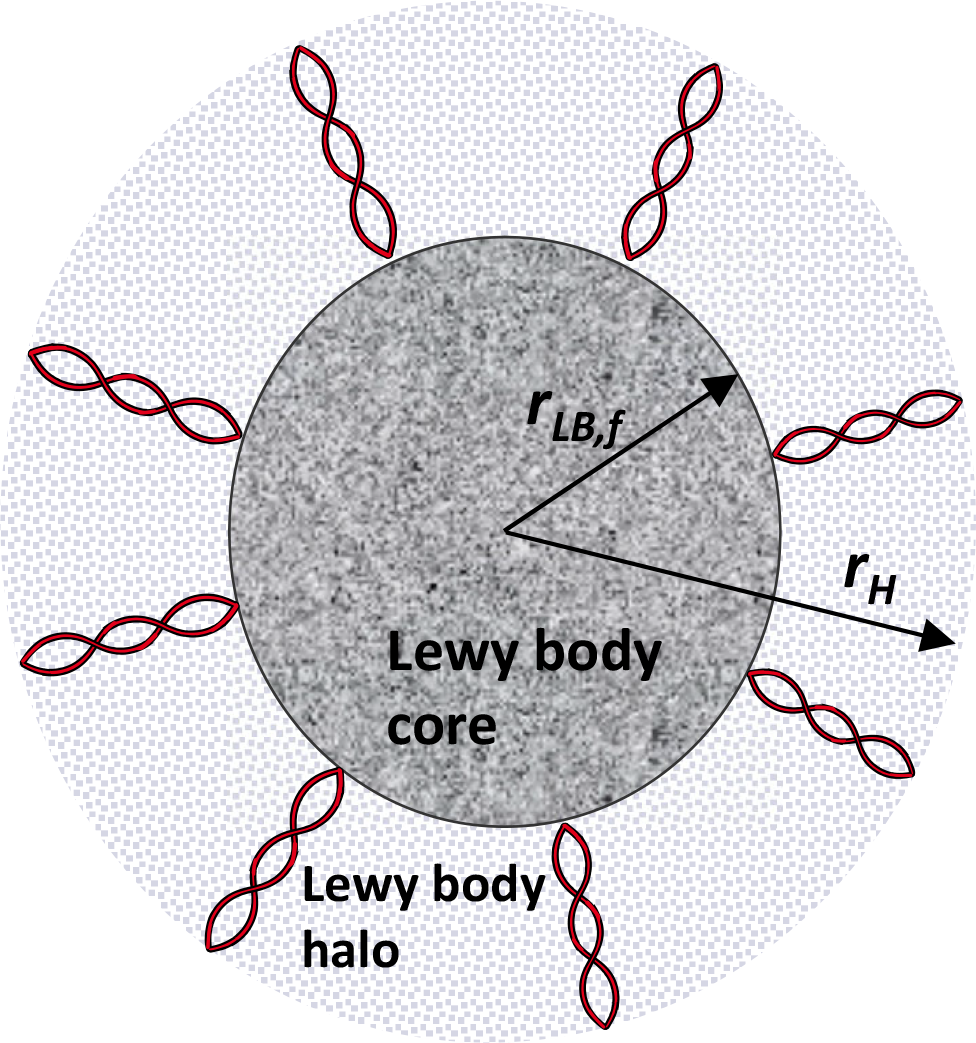
A diagram depicting an LB, which consists of two distinct regions: the central core made up of dense granular material and the surrounding halo containing radiating filaments (after electron micrographs presented in Fig. 1 in Olanow et al. (2004) and Fig. 3 in Fares et al. (2021)). *Figure generated with the aid of servier medical art, licensed under a creative commons attribution 3*.*0 generic license. http://Smart.servier.com*.

A minimal model for simulating LB growth, based on the aforementioned two-stage model, was developed in Kuznetsov and Kuznetsov (2022). Subsequent progress in modeling the growth of inclusions linked to neurodegenerative diseases was made in Kuznetsov (2023a), where the diffusion-driven transport of amyloid-β (Aβ) monomers between neurons was investigated. In Kuznetsov (2023b), a numerical study was conducted to explore the growth of Aβ plaques, a hallmark of Alzheimer’s disease. The present paper aims to provide a deeper understanding of the factors influencing LB growth and to formulate a simple analytical expression for LB radius versus time.

## 2. Materials and models

### 2.1. Governing equations

The model uses a single independent variable, which is time denoted as *t*. The model’s dependent variables are given in Table 1, while Table 2 provides a summary of the parameters used in the model.

**Table 1.** Dependent variables in the model.

| Symbol | Definition | Units |
| --- | --- | --- |
| $[A_{FR}]$ | Molar concentration of lipid membrane fragments, as well as damaged organelles, lysosomes, and impaired mitochondria – essentially, any fragments capable of forming the core of the LB – within the soma | $\mu\text{M}$ |
| $[B_{FR}]$ | Molar concentration of lipid membrane aggregates in the soma | $\mu\text{M}$ |
| $[A_{AS}]$ | Molar concentration of α-syn monomers in the soma | $\mu\text{M}$ |
| $[B_{AS}]$ | Molar concentration of α-syn fibrils in the soma | $\mu\text{M}$ |
| $r_{LB}$ | Radius of the growing core of the LB | $\mu\text{m}$ |
| $r_H$ | Radius of the growing halo | $\mu\text{m}$ |

**Table 2.**
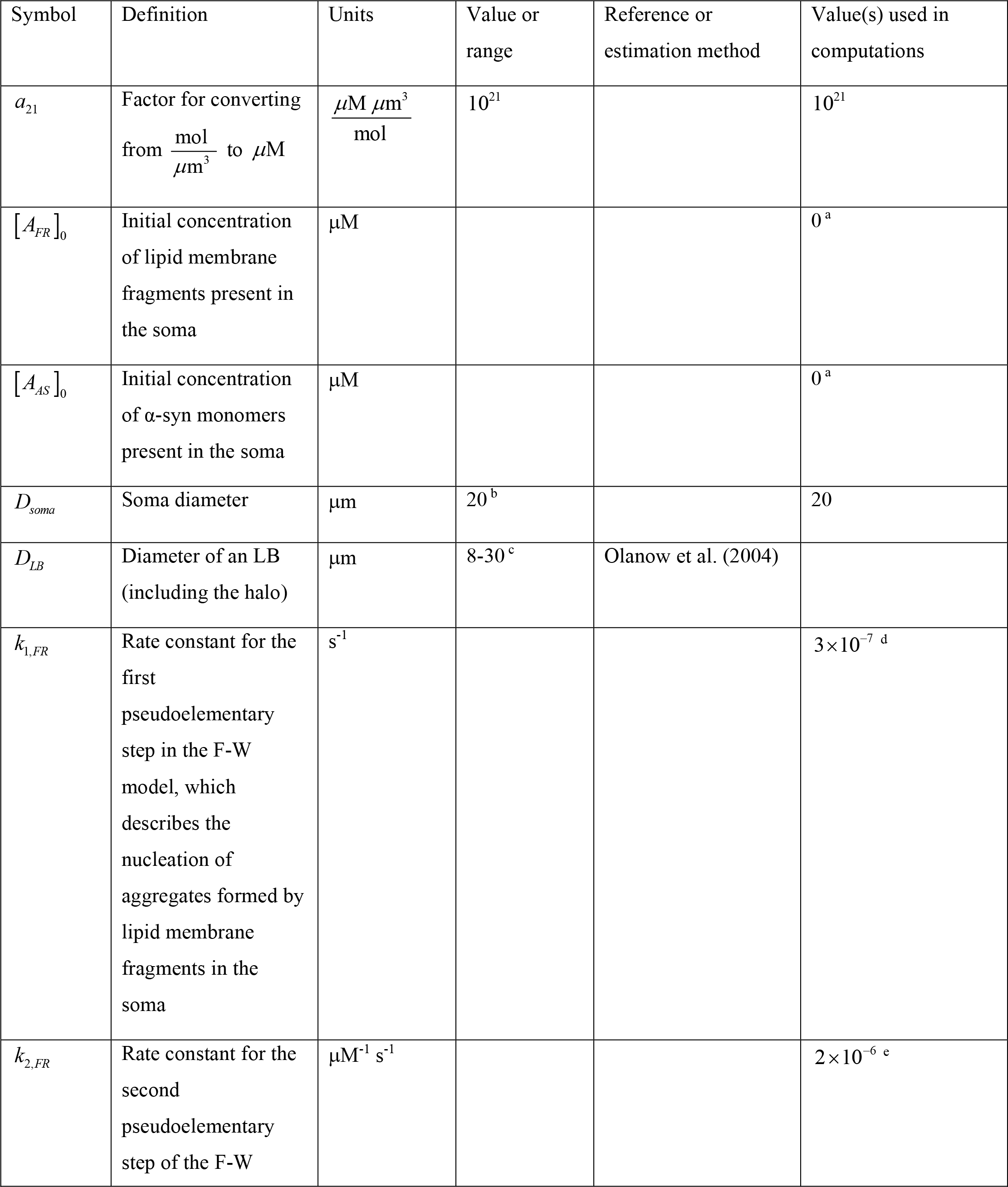
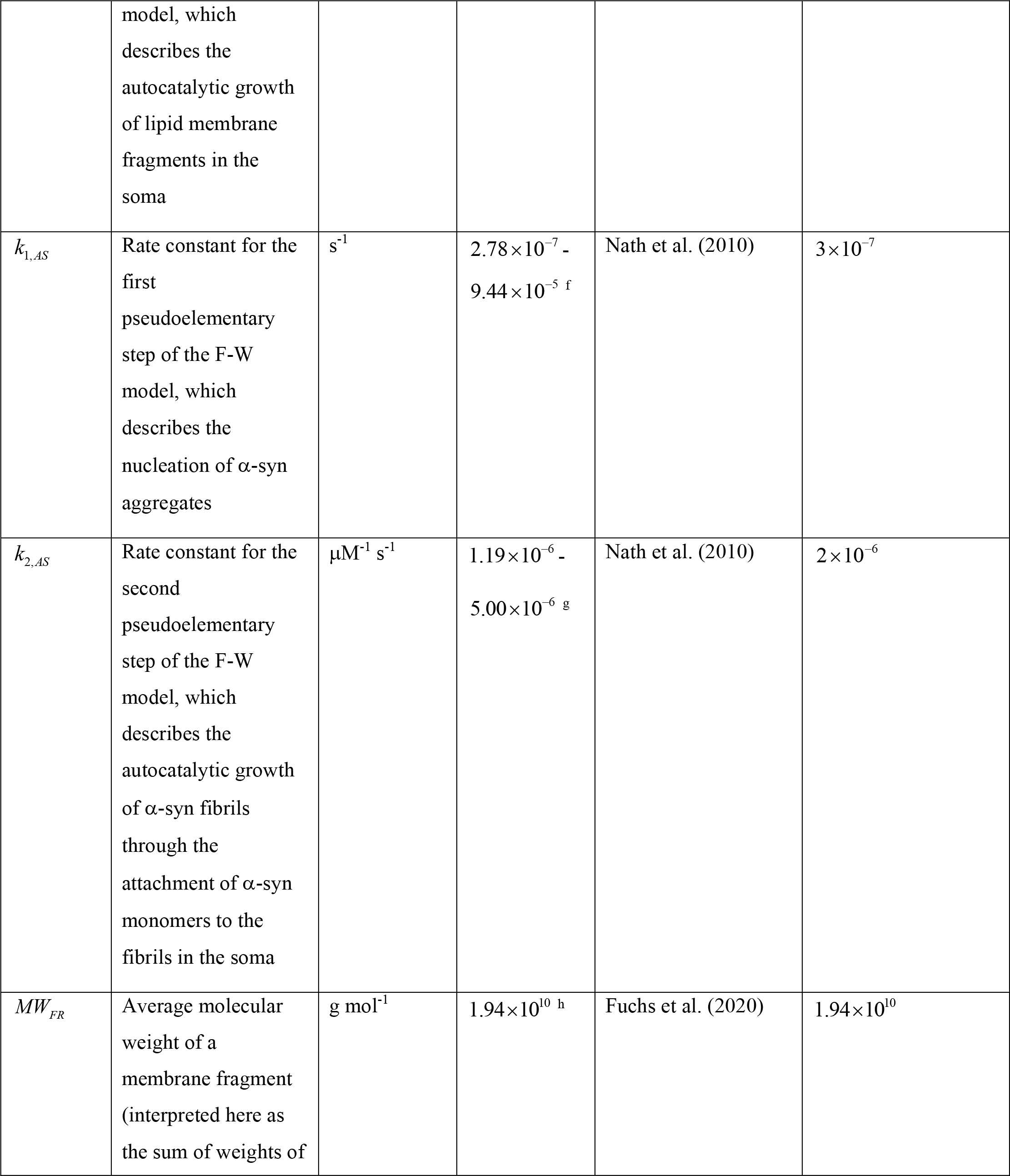
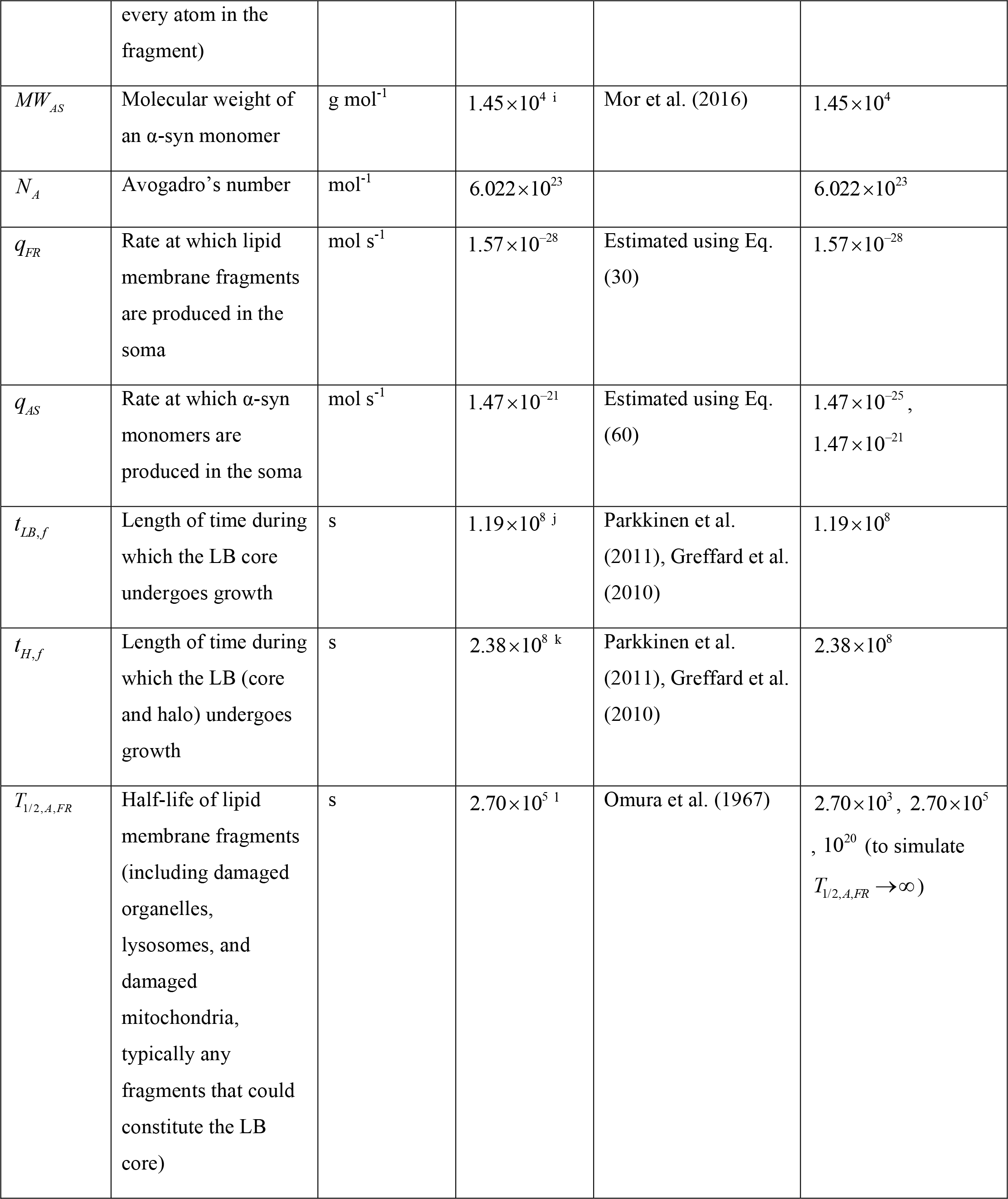
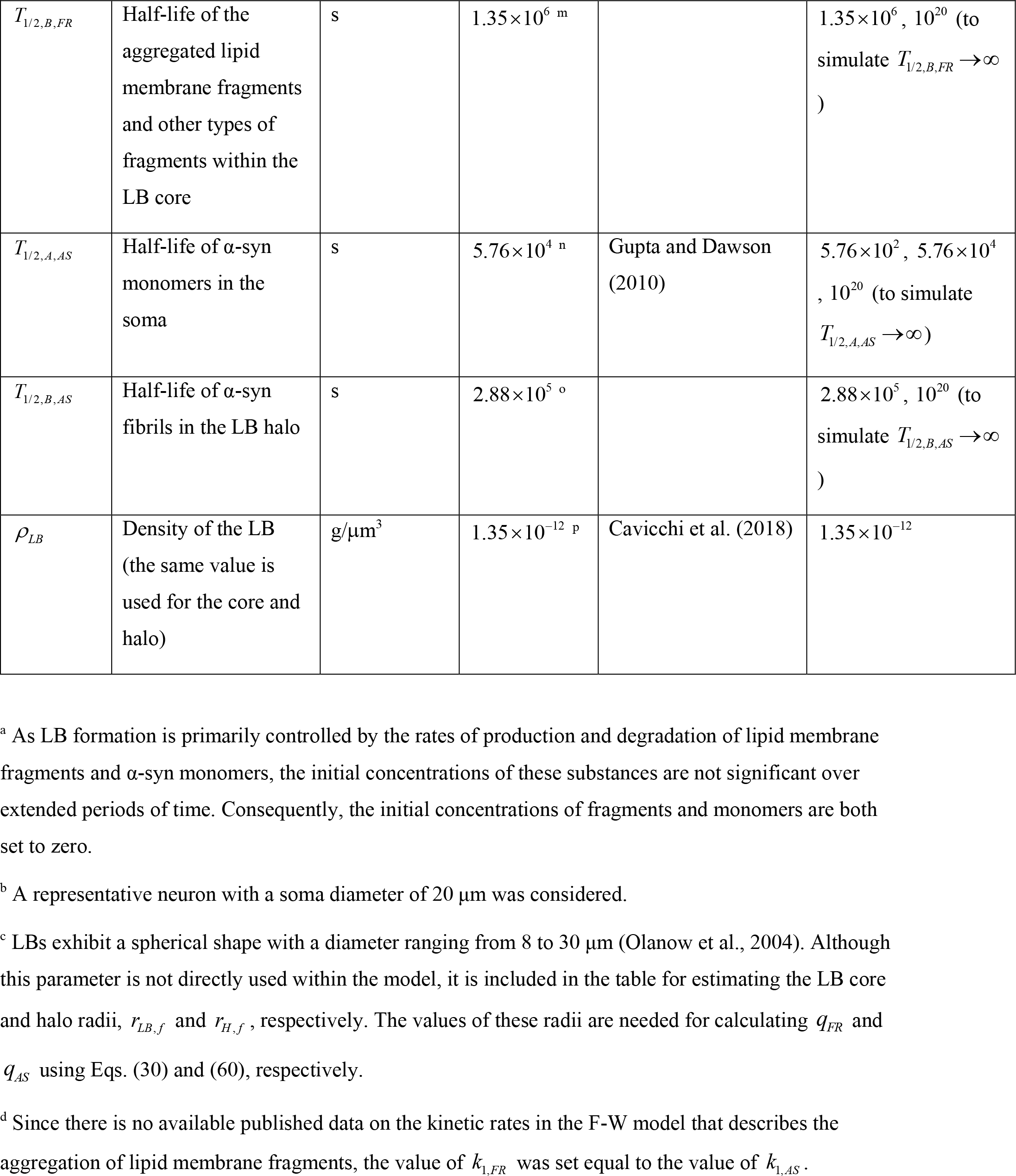
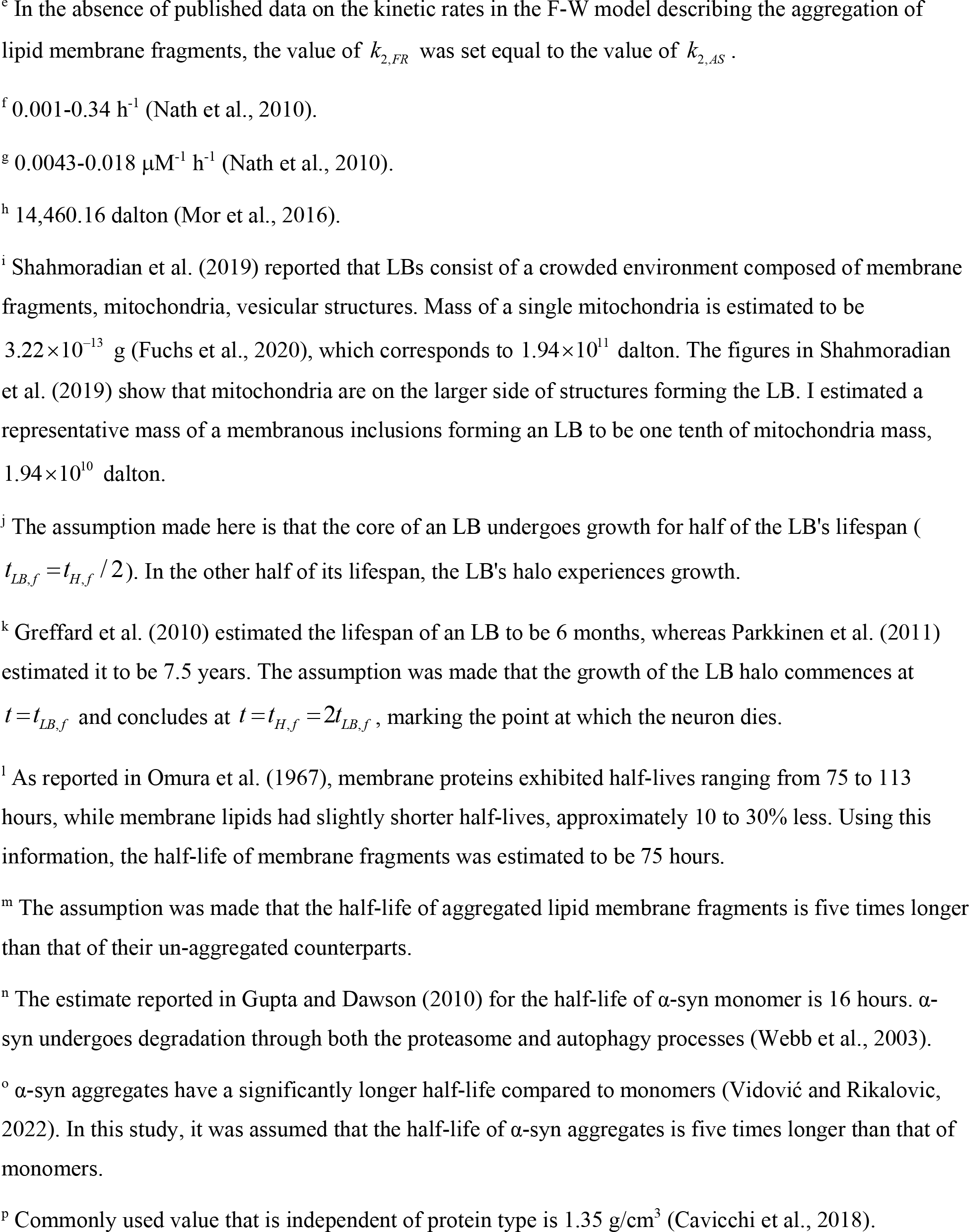
Model parameters and their estimated values.

| Symbol | Definition | Units | Value or range | Reference or estimation method | Value(s) used in computations |
| --- | --- | --- | --- | --- | --- |
| $a_{21}$ | Factor for converting from $\frac{\text{mol}}{\mu\text{m}^3}$ to $\mu\text{M}$ | $\frac{\mu\text{M } \mu\text{m}^3}{\text{mol}}$ | $10^{21}$ | | $10^{21}$ |
| $[A_{FR}]_0$ | Initial concentration of lipid membrane fragments present in the soma | $\mu\text{M}$ | | | $0^a$ |
| $[A_{AS}]_0$ | Initial concentration of α-syn monomers present in the soma | $\mu\text{M}$ | | | $0^a$ |
| $D_{soma}$ | Soma diameter | $\mu\text{m}$ | $20^b$ | | 20 |
| $D_{LB}$ | Diameter of an LB (including the halo) | $\mu\text{m}$ | $8\text{-}30^c$ | Olanow et al. (2004) | |
| $k_{1,FR}$ | Rate constant for the first pseudoelementary step in the F-W model, which describes the nucleation of aggregates formed by lipid membrane fragments in the soma | $\text{s}^{-1}$ | | | $3 \times 10^{-7}^d$ |
| $k_{2,FR}$ | Rate constant for the second pseudoelementary step of the F-W model, which describes the autocatalytic growth of lipid membrane fragments in the soma | $\mu\text{M}^{-1} \text{s}^{-1}$ | | | $2 \times 10^{-6}^e$ |
| $k_{1,AS}$ | Rate constant for the first pseudoelementary step of the F-W model, which describes the nucleation of α-syn aggregates | $s^{-1}$ | $2.78 \times 10^{-7}$ - $9.44 \times 10^{-5}$ <sup>f</sup> | Nath et al. (2010) | $3 \times 10^{-7}$ |
| $k_{2,AS}$ | Rate constant for the second pseudoelementary step of the F-W model, which describes the autocatalytic growth of α-syn fibrils through the attachment of α-syn monomers to the fibrils in the soma | $\mu M^{-1} s^{-1}$ | $1.19 \times 10^{-6}$ - $5.00 \times 10^{-6}$ <sup>g</sup> | Nath et al. (2010) | $2 \times 10^{-6}$ |
| $MW_{FR}$ | Average molecular weight of a membrane fragment (interpreted here as the sum of weights of every atom in the fragment) | $g\ mol^{-1}$ | $1.94 \times 10^{10}$ <sup>h</sup> | Fuchs et al. (2020) | $1.94 \times 10^{10}$ |
| $MW_{AS}$ | Molecular weight of an α-syn monomer | $\text{g mol}^{-1}$ | $1.45 \times 10^4$ <sup>i</sup> | Mor et al. (2016) | $1.45 \times 10^4$ |
| $N_A$ | Avogadro's number | $\text{mol}^{-1}$ | $6.022 \times 10^{23}$ | | $6.022 \times 10^{23}$ |
| $q_{FR}$ | Rate at which lipid membrane fragments are produced in the soma | $\text{mol s}^{-1}$ | $1.57 \times 10^{-28}$ | Estimated using Eq. (30) | $1.57 \times 10^{-28}$ |
| $q_{AS}$ | Rate at which α-syn monomers are produced in the soma | $\text{mol s}^{-1}$ | $1.47 \times 10^{-21}$ | Estimated using Eq. (60) | $1.47 \times 10^{-25}$,<br>$1.47 \times 10^{-21}$ |
| $t_{LB,f}$ | Length of time during which the LB core undergoes growth | s | $1.19 \times 10^8$ <sup>j</sup> | Parkkinen et al. (2011), Greffard et al. (2010) | $1.19 \times 10^8$ |
| $t_{H,f}$ | Length of time during which the LB (core and halo) undergoes growth | s | $2.38 \times 10^8$ <sup>k</sup> | Parkkinen et al. (2011), Greffard et al. (2010) | $2.38 \times 10^8$ |
| $T_{1/2,A,FR}$ | Half-life of lipid membrane fragments (including damaged organelles, lysosomes, and damaged mitochondria, typically any fragments that could constitute the LB core) | s | $2.70 \times 10^5$ <sup>l</sup> | Omura et al. (1967) | $2.70 \times 10^3$, $2.70 \times 10^5$, $10^{20}$ (to simulate $T_{1/2,A,FR} \rightarrow \infty$) |
| $T_{1/2,B,FR}$ | Half-life of the aggregated lipid membrane fragments and other types of fragments within the LB core | s | $1.35 \times 10^6$ <sup>m</sup> | | $1.35 \times 10^6, 10^{20}$ (to simulate $T_{1/2,B,FR} \rightarrow \infty$) |
| $T_{1/2,A,AS}$ | Half-life of α-syn monomers in the soma | s | $5.76 \times 10^4$ <sup>n</sup> | Gupta and Dawson (2010) | $5.76 \times 10^2, 5.76 \times 10^4, 10^{20}$ (to simulate $T_{1/2,A,AS} \rightarrow \infty$) |
| $T_{1/2,B,AS}$ | Half-life of α-syn fibrils in the LB halo | s | $2.88 \times 10^5$ <sup>o</sup> | | $2.88 \times 10^5, 10^{20}$ (to simulate $T_{1/2,B,AS} \rightarrow \infty$) |
| $\rho_{LB}$ | Density of the LB (the same value is used for the core and halo) | $\text{g}/\mu\text{m}^3$ | $1.35 \times 10^{-12}$ <sup>p</sup> | Cavicchi et al. (2018) | $1.35 \times 10^{-12}$ |

<sup>a</sup> As LB formation is primarily controlled by the rates of production and degradation of lipid membrane fragments and α-syn monomers, the initial concentrations of these substances are not significant over extended periods of time. Consequently, the initial concentrations of fragments and monomers are both set to zero.
<sup>b</sup> A representative neuron with a soma diameter of 20 $\mu\text{m}$ was considered.
<sup>c</sup> LBs exhibit a spherical shape with a diameter ranging from 8 to 30 $\mu\text{m}$ (Olanow et al., 2004). Although this parameter is not directly used within the model, it is included in the table for estimating the LB core and halo radii, $r_{LB,f}$ and $r_{H,f}$ , respectively. The values of these radii are needed for calculating $q_{FR}$ and $q_{AS}$ using Eqs. (30) and (60), respectively.
<sup>d</sup> Since there is no available published data on the kinetic rates in the F-W model that describes the aggregation of lipid membrane fragments, the value of $k_{1,FR}$ was set equal to the value of $k_{1,AS}$ .
<sup>e</sup> In the absence of published data on the kinetic rates in the F-W model describing the aggregation of lipid membrane fragments, the value of $k_{2,FR}$ was set equal to the value of $k_{2,AS}$ .
<sup>f</sup> 0.001-0.34 h<sup>-1</sup> (Nath et al., 2010).
<sup>g</sup> 0.0043-0.018 μM<sup>-1</sup> h<sup>-1</sup> (Nath et al., 2010).
<sup>h</sup> 14,460.16 dalton (Mor et al., 2016).
<sup>i</sup> Shahmoradian et al. (2019) reported that LBs consist of a crowded environment composed of membrane fragments, mitochondria, vesicular structures. Mass of a single mitochondria is estimated to be $3.22 \times 10^{-13}$ g (Fuchs et al., 2020), which corresponds to $1.94 \times 10^{11}$ dalton. The figures in Shahmoradian et al. (2019) show that mitochondria are on the larger side of structures forming the LB. I estimated a representative mass of a membranous inclusions forming an LB to be one tenth of mitochondria mass, $1.94 \times 10^{10}$ dalton.
<sup>j</sup> The assumption made here is that the core of an LB undergoes growth for half of the LB's lifespan ( $t_{LB,f} = t_{H,f} / 2$ ). In the other half of its lifespan, the LB's halo experiences growth.
<sup>k</sup> Greffard et al. (2010) estimated the lifespan of an LB to be 6 months, whereas Parkkinen et al. (2011) estimated it to be 7.5 years. The assumption was made that the growth of the LB halo commences at $t = t_{LB,f}$ and concludes at $t = t_{H,f} = 2t_{LB,f}$ , marking the point at which the neuron dies.
<sup>l</sup> As reported in Omura et al. (1967), membrane proteins exhibited half-lives ranging from 75 to 113 hours, while membrane lipids had slightly shorter half-lives, approximately 10 to 30% less. Using this information, the half-life of membrane fragments was estimated to be 75 hours.
<sup>m</sup> The assumption was made that the half-life of aggregated lipid membrane fragments is five times longer than that of their un-aggregated counterparts.
<sup>n</sup> The estimate reported in Gupta and Dawson (2010) for the half-life of α-syn monomer is 16 hours. α-syn undergoes degradation through both the proteasome and autophagy processes (Webb et al., 2003).

In order to model the formation of lipid fragment aggregates in the soma, a phenomenological two-step Finke-Watzky (F-W) model, as described in Morris et al. (2008), Iashchishyn et al. (2017), was used. This model simulates polymer aggregation by employing two parallel, competing processes: continuous nucleation and autocatalytic surface growth (Finke et al., 2020). The F-W model has previously been applied to fit data related to the aggregation of neurological proteins (Morris et al., 2008). The present model does not simulate the process of how sticky membrane fragment aggregates assemble into an LB core. It is assumed that this assembly occurs more rapidly than the formation of aggregates in the soma. The F-W model can be delineated through the following transitions:

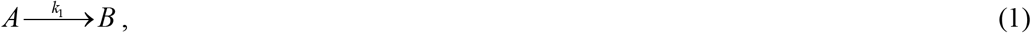

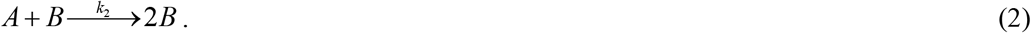

Typically, the F-W model is used to simulate the conversion of an initial concentration of monomers, denoted as [*A*]_0_, into a misfolded or active (capable of autocatalysis) form *B* (Iashchishyn et al., 2017). The active form *B* is commonly linked to aggregates. Analytical solutions for such scenarios have been documented (see, for example, Bentea et al. (2017)). Here the F-W model is applied to a different scenario. Recent findings reported in Shahmoradian et al. (2019) and the analysis presented in Fares et al. (2021) propose that the core of the LB comprises lipid membrane fragments and damaged organelles. For conciseness, these fragmented or damaged components are referred to as membrane fragments.

It is assumed that membrane fragments are generated at a constant rate, *q*_*FR*_, within the soma. Stating the conservation of membrane fragments in the soma leads to the following equation:

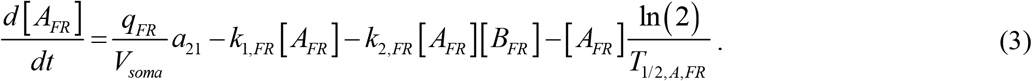

In this equation, 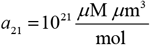 represents the conversion factor from 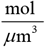 to *μ*M. The first term on the right-hand side of Eq. (3) accounts for the rate at which membrane fragments are produced. The second term models the rate of conversion of membrane fragments into aggregates through nucleation, while the third term represents the rate of conversion of membrane fragments into aggregates through autocatalytic growth. The second and third terms are formulated by applying the law of mass action to describe the rates of reactions presented in Eqs. (1) and (2). The final term on the right-hand side Eq. (3) simulates the degradation of membrane fragments.

Expressing the conservation of membrane aggregates within the soma, which constitute the core of the LB, results in the following equation:

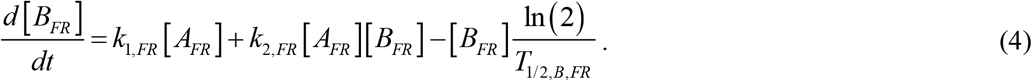

On the right-hand side of Eq. (4), the first term accounts for the rate at which aggregates are produced by nucleation, while the second term represents the rate at which aggregates are produced through autocatalytic growth. These terms are equal in magnitude but have opposite signs compared to the second and third terms on the right-hand side of Eq. (3). This is because, in the F-W model, the rate of aggregate production is equivalent to the rate of monomer consumption. The final term on the right-hand side of Eq. (4) models the degradation of lipid membrane aggregates.

Eqs. (3) and (4) are solved subject to the following initial conditions:

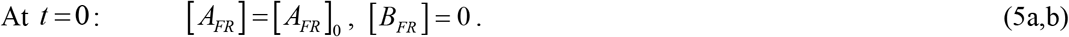

The dimensionless form of Eqs. (3) and (4) is as follows:

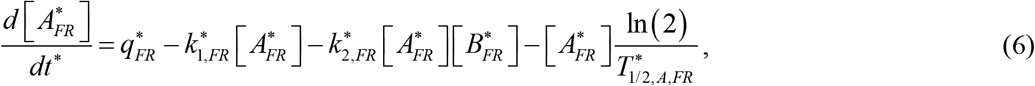

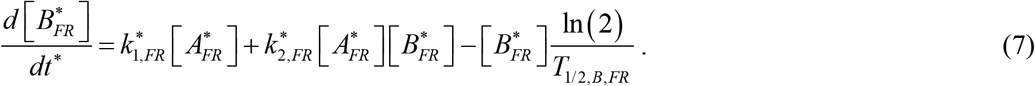

The dimensionless form of the initial conditions given by Eq. (5) can be expressed as follows:

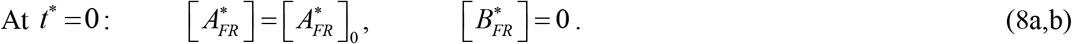

Table 3 defines the dimensionless independent variable, while Table 4 summarizes dimensionless dependent variables. Table 5 gives the dimensionless parameters utilized in the model.

**Table 3.**
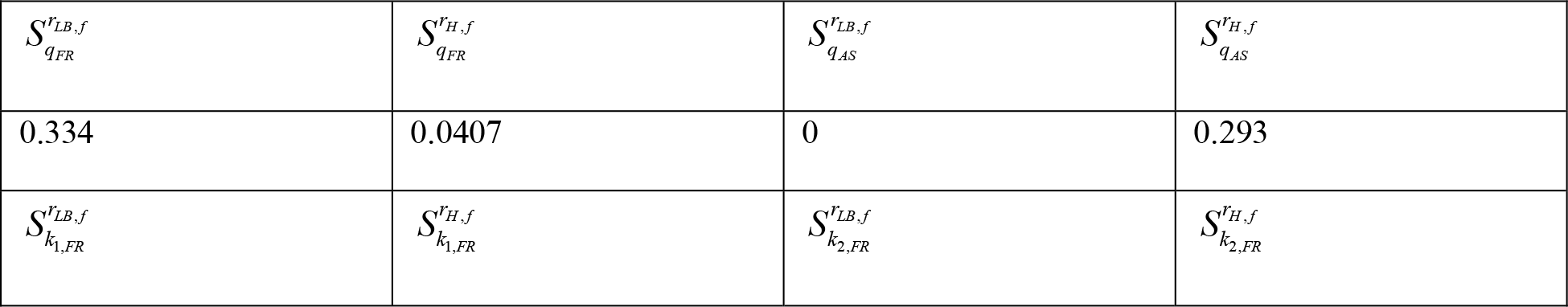
Independent variable in dimensionless form in the model.

**Table 4.** Dimensionless dependent variables used in the model.

| Symbol | Definition |
| --- | --- |
| $[A_{FR}^*]$ | $[A_{FR}] k_{2,AS} / k_{1,AS}$ |
| $[B_{FR}^*]$ | $[B_{FR}] k_{2,AS} / k_{1,AS}$ |
| $[A_{AS}^*]$ | $[A_{AS}] k_{2,AS} / k_{1,AS}$ |
| $[B_{AS}^*]$ | $[B_{AS}] k_{2,AS} / k_{1,AS}$ |
| $r_{LB}^*$ | $r_{LB} / D_{soma}$ |
| $r_H^*$ | $r_H / D_{soma}$ |

**Table 5.**
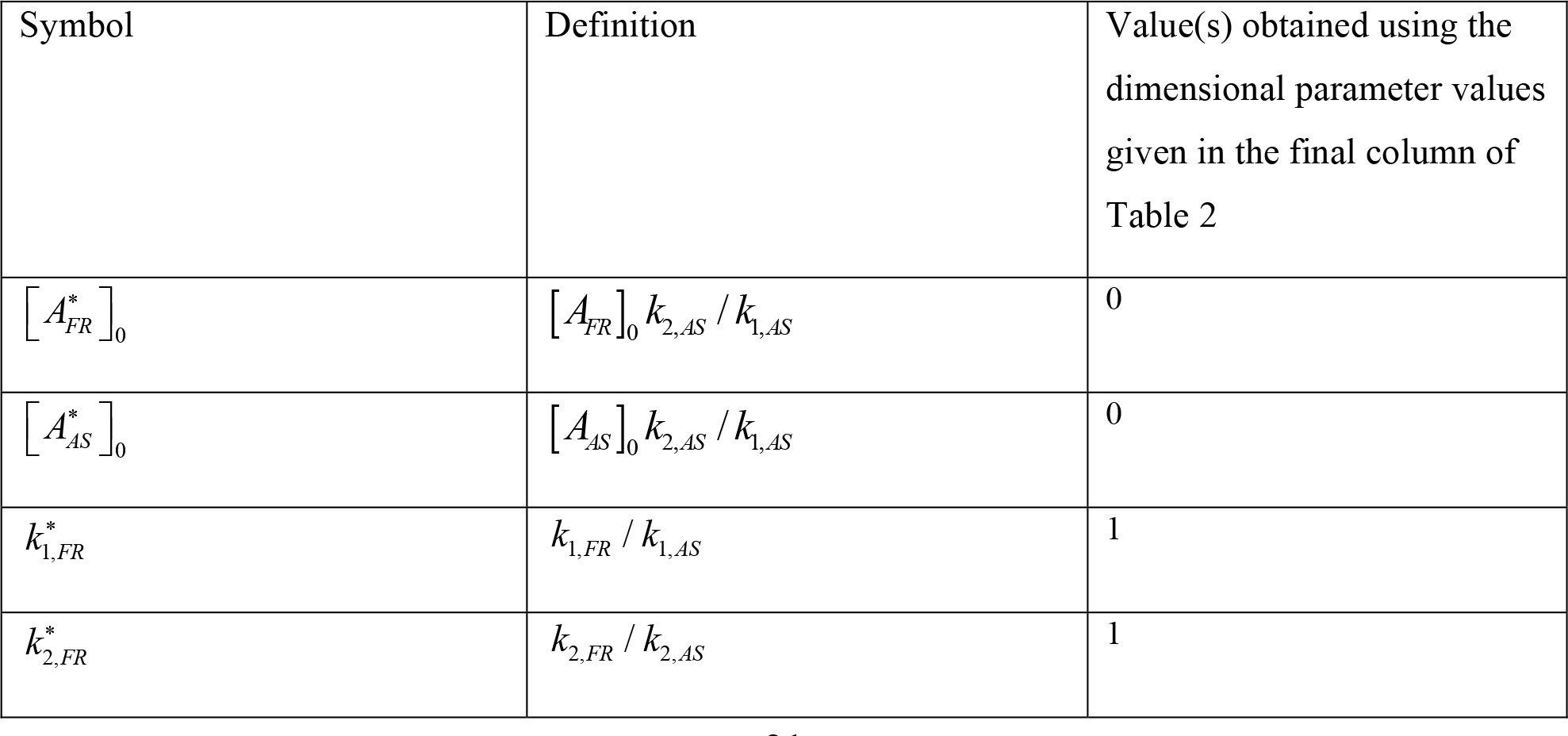
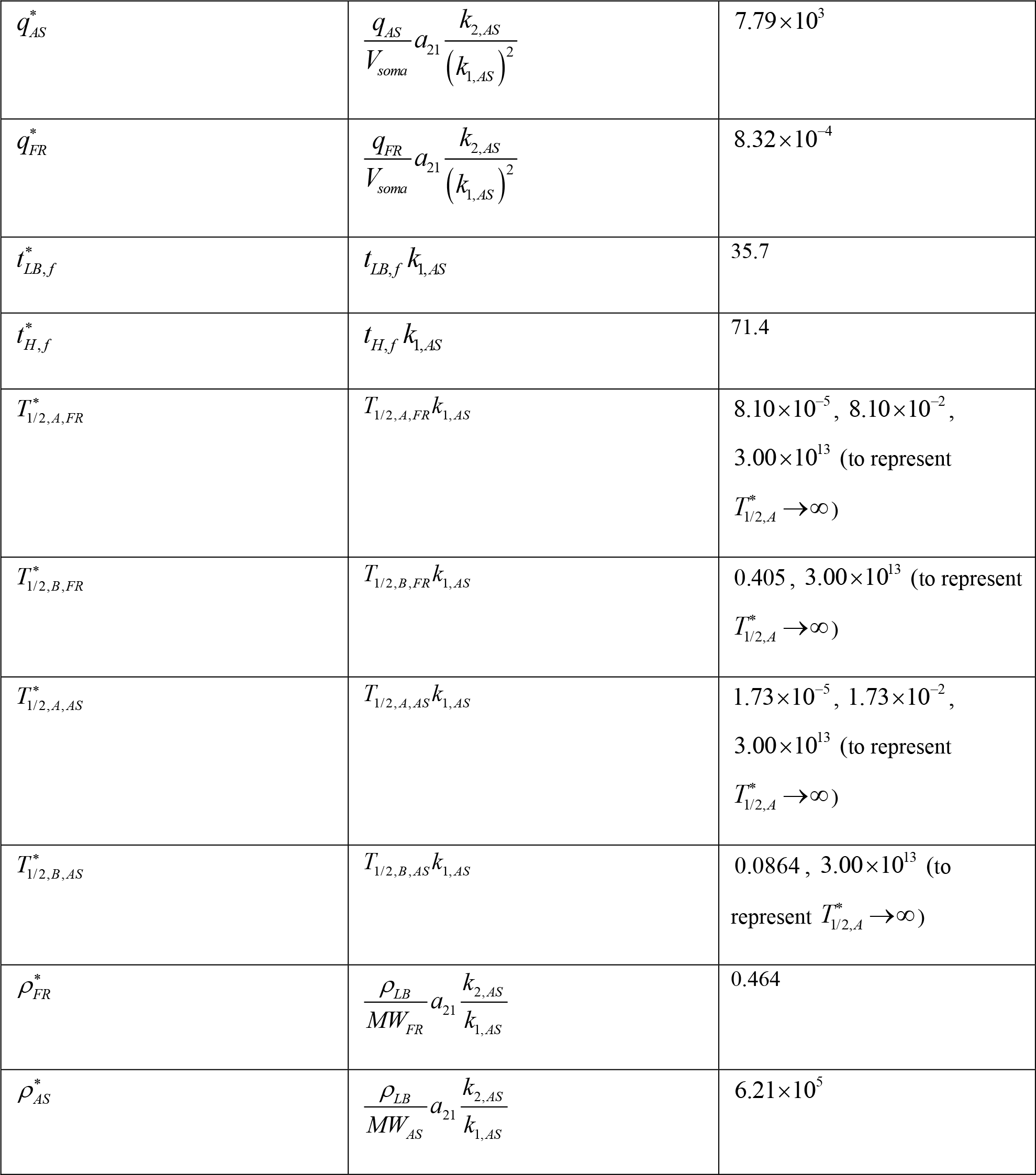
Dimensionless parameters used in the model.

When Eqs. (6) and (7) were added together, the following result was obtained:

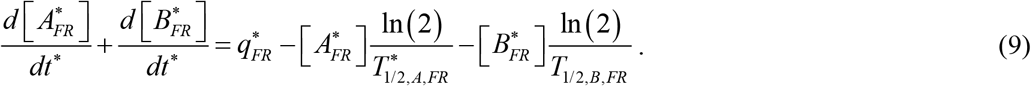

By integrating Eq. (9) with respect to time under the conditions of 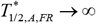 and 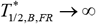, and utilizing the initial condition provided in Eq. (8), the following was obtained:

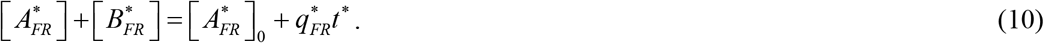

The increase 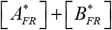 over time is attributed to the continuous production of lipid membrane fragments in the soma.

By eliminating 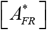 from Eq. (7) using Eq. (10), the following equation was obtained:

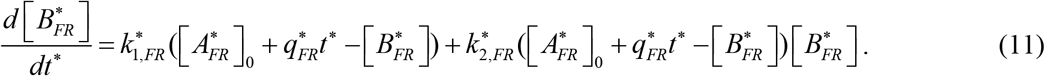

Eq. (11) is similar to the one analyzed in Kuznetsov and Kuznetsov (2022). To determine the exact analytical solution of Eq. (11) with the initial condition given by Eq. (8b), the DSolve function followed by the FullSimplify function in Mathematica 13.3 (Wolfram Research, Champaign, IL) was utilized. The obtained solution is presented below:

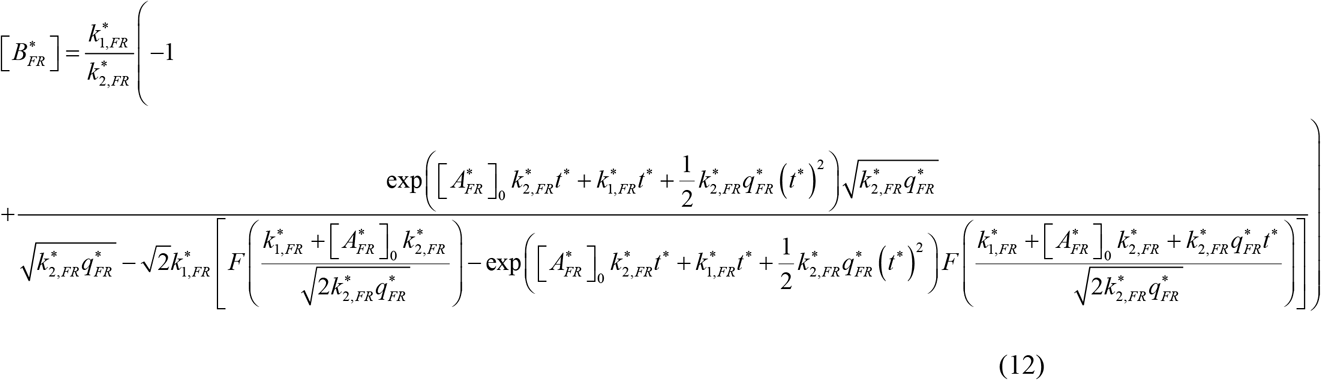

In this equation, the symbol *F* (*x*) represents Dawson’s integral:

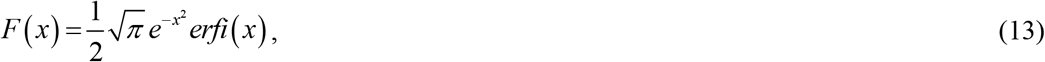

where *erfi* (*x*) is the imaginary error function.

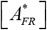 can then be obtained using Eq. (10) as follows:

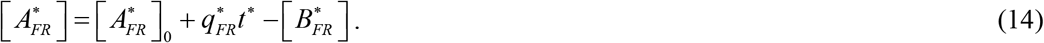

The exact solution provided by Eqs. (12) and (14) is rather complex. However, a more elegant approximate solution, applicable when 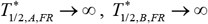, and 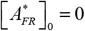, can be derived as follows. Numerical analysis of Eqs. (6)-(8) reveals that 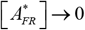 when *t*^*^ →∞. By substituting this into Eq. (10), the following is obtained:

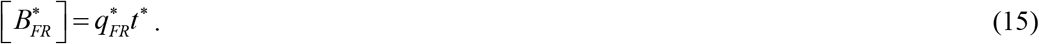

Reverting to dimensional variables, Eq. (15) can be expressed as follows:

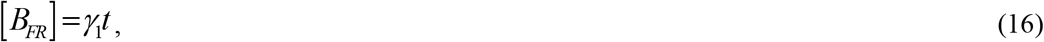

where

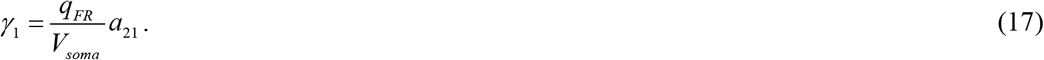

Substituting Eq. (15) into Eq. (7) leads to:

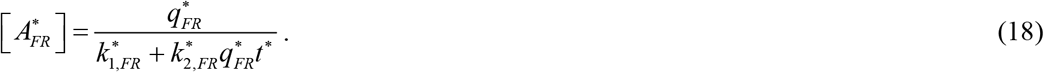

Going back to dimensional variables, Eq. (18) can be rewritten as follows:

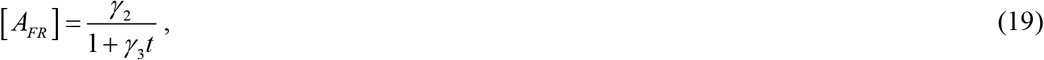

where

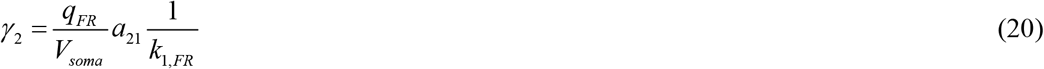

and

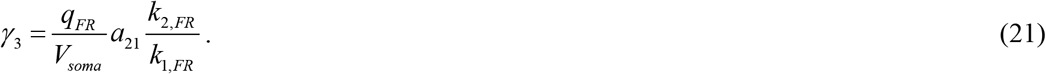

It is essential to note that Eqs. (15) and (18) are useful for extended periods. However, they are not applicable when *t*^*^ → 0; for example, Eq. (18) results in 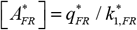, which contradicts the initial condition defined in Eq. (8a).

The growth of the LB’s core (Fig. 1) is calculated as follows. The total number of membrane fragments incorporated into the LB core at any time *t* is calculated as follows (Watzky et al., 2008):

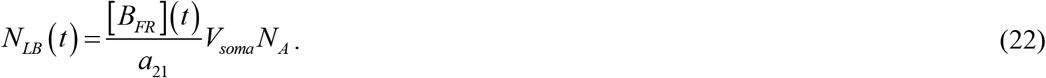

Alternatively, *N*_*LB*_ (*t*) can as calculated as (Watzky et al., 2008):

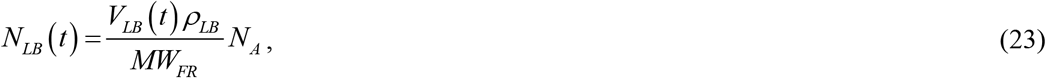

where *MW*_*FR*_ represents the average molecular weight of a membrane fragment, which is the sum of the weights of all the atoms within the fragment.

By setting the expressions on the right-hand sides of Eqs. (22) and (23) equal to each other and solving for the volume of the LB, the following result is obtained:

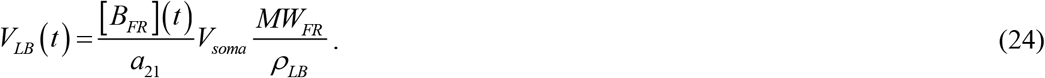

Assuming that the LB core is of spherical shape,

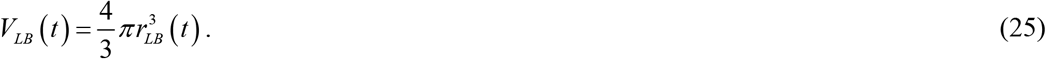

When Eqs. (24) and (25) are solved for the radius of the LB core, the following equation is obtained:

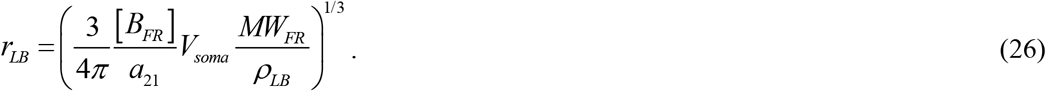

In dimensionless variables, Eq. (26) can be expressed as follows:

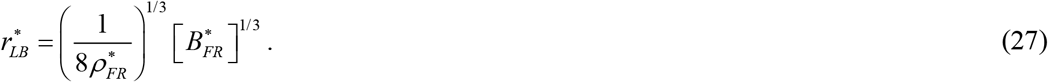

This is consistent with findings of Szabo and Lente (2019), which noted that the volume of a nanoparticle is directly proportional to the number of monomers it comprises. Consequently, the size of a nanoparticle increases in proportion to the cube root of the number of monomers forming the particle.

By substituting Eq. (15) into Eq. (27), the following equation for the dimensionless radius of the LB core is derived:

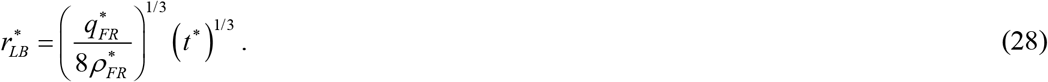

Returning back to the dimensional variables, the following equation is obtained:

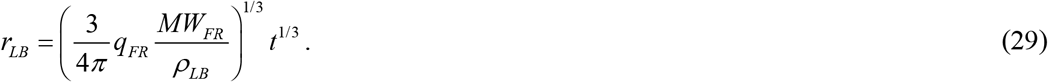

The growth of the LB’s core continues until *t* reaches *t*_*LB, f*_, at which point *r*_*LB*_ reaches *r*_*LB, f*_. Solving Eq. (29) for this case, the following is obtained:

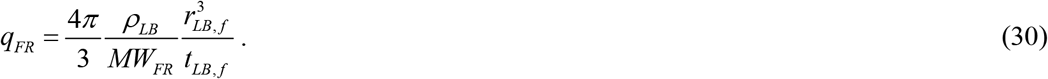

Eq. (30) plays a crucial role in the estimation of model parameters as it enables the calculation of *q*_*FR*_ when *r*_*LB, f*_ and *t*_*LB, f*_ are known. Eq. (30) was utilized to obtain an estimate for *q*_*FR*_ of 1.57 × 10^−28^ mol s^-1^, which is reported in Table 2.

Eq. (29) can be reformulated as follows:

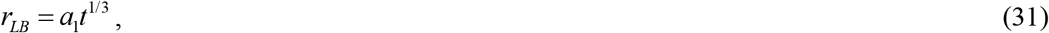

where

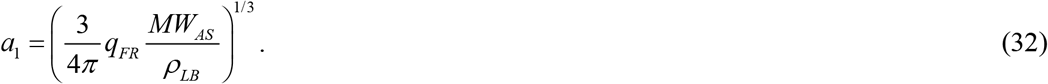

Eq. (31) indicates that the size of the LB core increases in direct proportion to the cube root of time. Interestingly, Eqs. (31) and (32) reveal that the radius of an LB core remains unaffected by the kinetic constants governing the aggregation rate of lipid membrane fragments, *k*_1, *FR*_ and *k*_2,*FR*_. This suggests that the kinetics of lipid membrane fragment aggregation is not a limiting factor in the aggregation process. Instead, the growth rate of the LB core is constrained by the rate at which lipid membrane fragments are produced and how quickly these fragments degrade. Eq. (31) can be used for fitting experimental data in future studies.

An assumption was made that α-syn monomers are produced within the soma at a rate denoted as *q*_*AS*_. The F-W model, as represented by Eqs. (3) and (4), was once again utilized, this time to simulate the formation of α-syn fibrils (Morris et al., 2008) within the soma. It is important to note that the model developed here does not account for the process of how adhesive α-syn filaments assemble into an LB halo (Cookson, 2005). The assumption made here is that this assembly process occurs more rapidly than the formation of filaments within the soma. Expressing the conservation of α-syn monomers in the soma leads to the following equation:

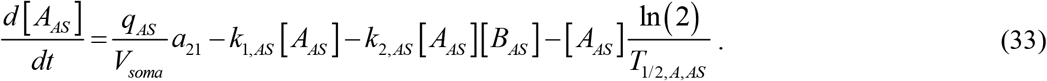

Stating the conservation of α-syn fibrils, which form the halo region of the LB, the following equation was obtained:

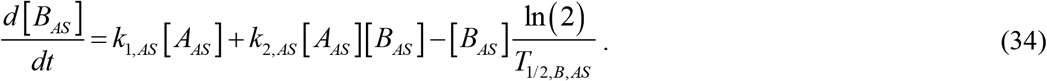

Eqs. (33) and (34) are similar to Eqs. (3) and (4). The first term on the right-hand side of Eq. (33) represents the rate at which α-syn monomers are generated. The second and third terms model the conversion of α-syn monomers into fibrils through nucleation and autocatalytic growth processes, respectively. The fourth term accounts for the decay of α-syn monomers due to their limited half-life. The two terms on the right-hand side of Eq. (34) represent the rate of α-syn fibril concentration increase attributed to nucleation and autocatalytic growth, respectively, while the third term simulates degradation of α-syn fibrils due to their finite half-life.

Eqs (33) and (34) are solved with the following initial condition:

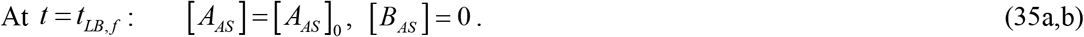

The dimensionless form of Eqs. (33) and (34) is as follows:

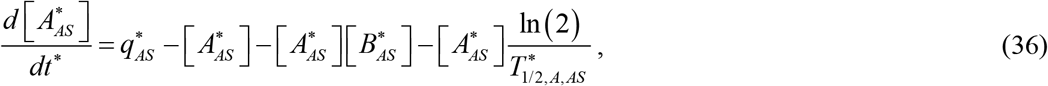

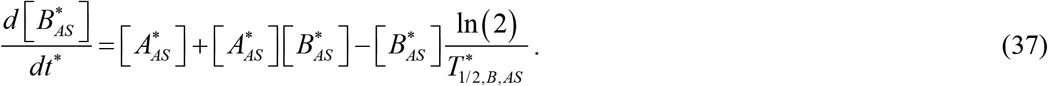

The dimensionless form of the initial conditions described by Eq. (35) can be expressed as follows:

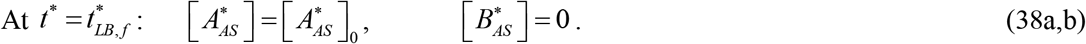

The initial value problem defined by Eqs. (36)-(38) becomes analogous to that presented by Eqs. (6)-(8) when the following transformation is utilized:

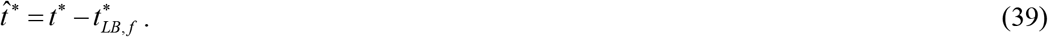

By adding Eqs. (36) and (37) and integrating the result with respect to 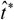 under the constraints of 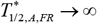 and 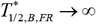, the following equation is obtained:

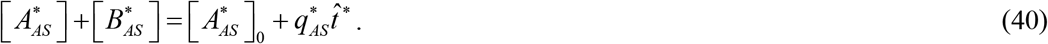

The increase of 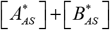 with time is attributed to the production of α-syn monomers in the soma.

By eliminating 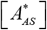 from Eq. (37) using Eq. (40), the following result is obtained:

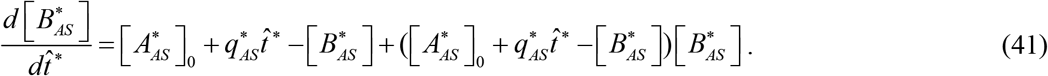

The exact solution to Eq. (41) with the initial condition (38b) is as follows:

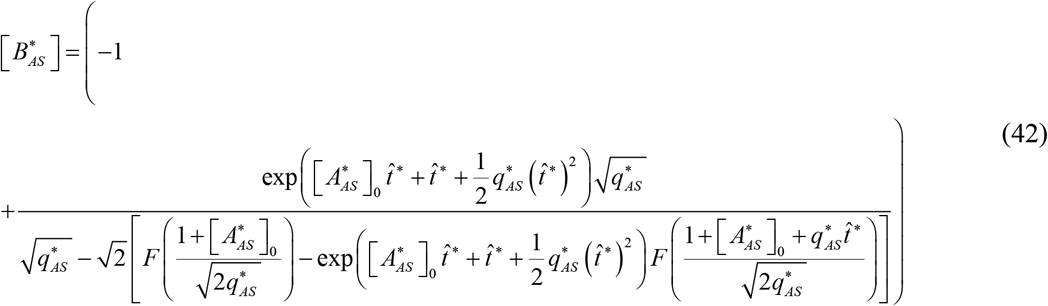

We can determine 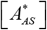 using Eq. (40) as follows:

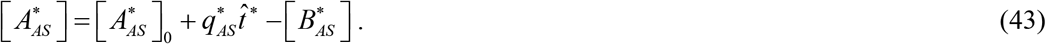

The exact solution given by Eqs. (42) and (43) is complex. A simpler approximate solution can be obtained for 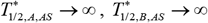, and 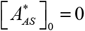. By numerically solving Eqs. (36)-(38), we found that 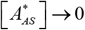 when *t*^*^ →∞. Substituting this into Eq. (40) leads to:

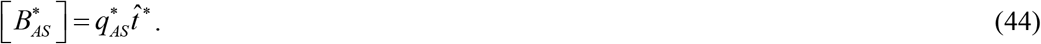

When reverting to the dimensional variables, the following solution for [B_*AS*_] is obtained:

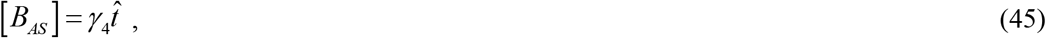

where

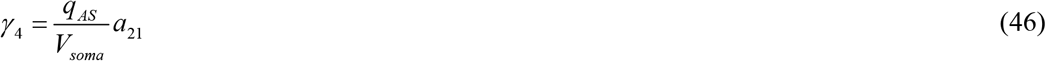

and

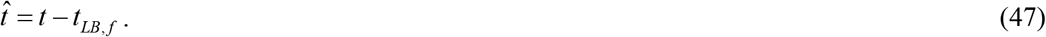

Substituting Eq. (44) into Eq. (37) results in the following:

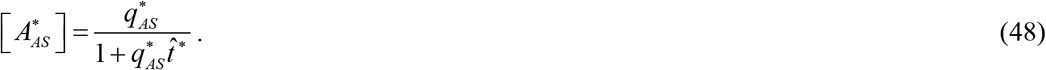

Switching back to dimensionless variables, the following is obtained:

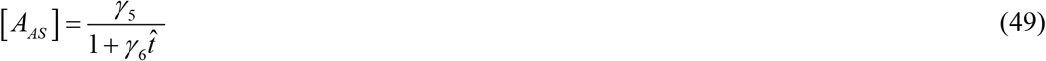

where

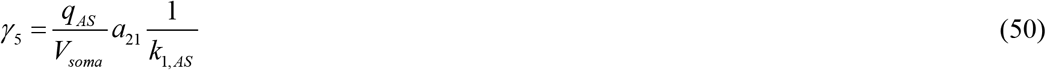

and

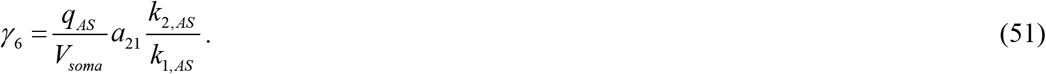

Eqs. (44) and (48) are useful only when 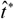 is large. However, they are not applicable when 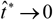. For instance, at 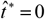 Eq. (48) yields 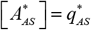, which contradicts the initial condition specified in Eq. (38a).

To compute the growth of the halo surrounding the LB core (Fig. 1) the following method was utilized. The total number of α-syn monomers incorporated into the LB halo was calculated as follows (Watzky et al., 2008):

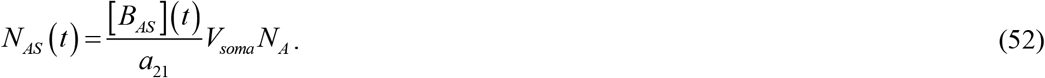

On the other hand *N*_*AS*_ (*t*) can be determined from the following equation (Watzky et al., 2008):

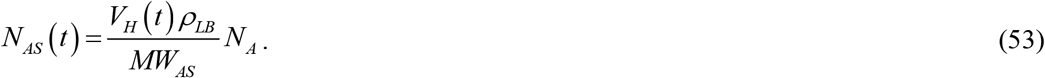

Here, *MW*_*AS*_ stands for the molecular weight of an α-syn monomer, while *V*_*H*_ represents the volume of the LB’s halo.

Equating the right-hand sides of Eqs. (52) and (53) and solving for the LB volume, the following is obtained:

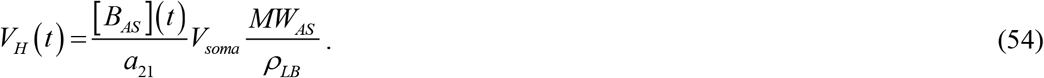

It is assumed that the LB halo takes the form of a spherical shell,

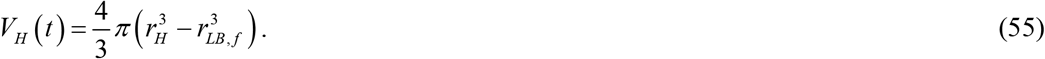

Solving Eqs. (54) and (55) for the radius of the halo, the following result is obtained:

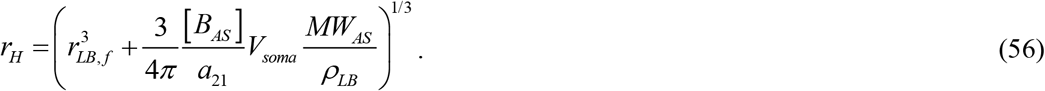

When using dimensionless variables, Eq. (56) can be expressed as follows:

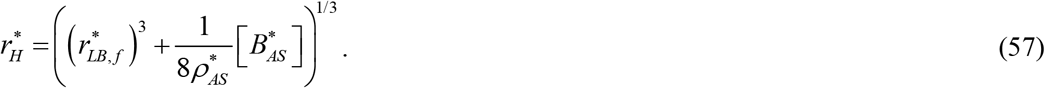

Substituting the approximate solution (44) into Eq. (57) yields the following result:

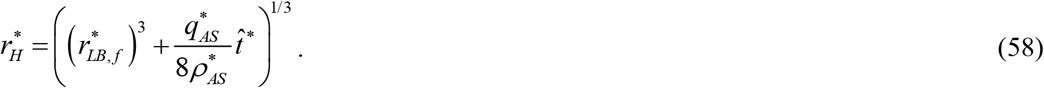

Returning to dimensional variables, the following is obtained:

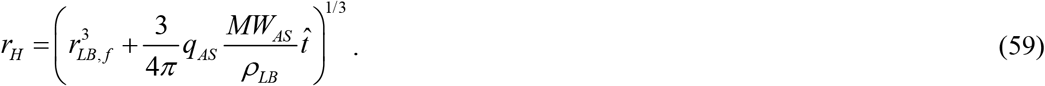

The growth of the LB’s halo continues until 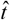 reaches 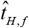, at which point 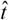 reaches 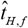. Solving Eq. (59) for *q*_*AS*_ in this particular scenario, the following result is obtained:

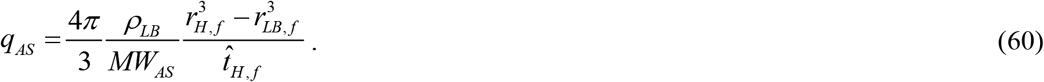

Eq. (60) plays an important role in estimating model parameters as it enables the calculation of *q*_*AS*_ when the values of *r*_*LB*,,*f*_, *r*_*H,f*_, and 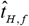 are known. Eq. (60) was used to estimate *q*_*AS*_ (a value of 1.47 ×10^−21^ mol s^-1^ reported in Table 2).

Eq. (59) can be reformulated as follows:

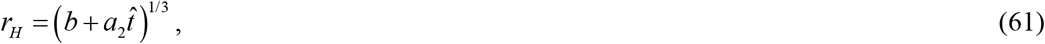

where

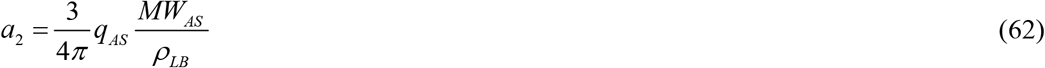

and

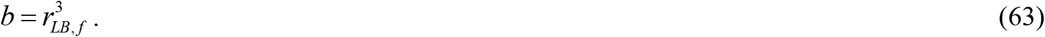

Eq. (61) can be employed for fitting future experimental data.

Table 3 gives the dimensionless independent variable, whereas Table 4 summarizes the dimensionless dependent variables. Table 5 sums up the dimensionless parameters employed within the model.

### 2.2. Sensitivity analysis

The dependences of radii of a fully grown LB core, *r*_*LB, f*_, and a fully grown LB halo, *r*_*H, f*_, on various model parameters were investigated. To conduct this analysis, the local sensitivity coefficients, representing the first-order partial derivatives of the core radius and halo radius with respect to various model parameters, were computed following the methods outlined in Beck and Arnold (1977), Zadeh and Montas (2010), Zi (2011), Kuznetsov and Kuznetsov (2019). To determine, for example, the sensitivity of *r*_*H,f*_ to *q*_*FR*_, the following procedure was employed:

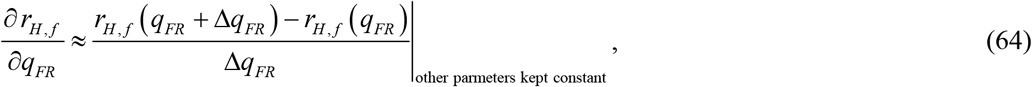

where Δ*q*_*FR*_ = 10^−3^ *q*_*FR*_ represents the step size. To confirm the sensitivity coefficients’ independence of the step size, various step sizes were tested.

To obtain quantities that are independent of units used to measure the parameters, non-dimensionalized relative sensitivity coefficients were computed, following the approach described in Zadeh and Montas (2010). For example:

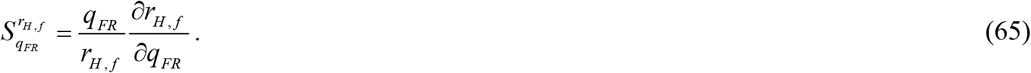

As per Kacser et al. (1995), the parameter defined in Eq. (65) can be interpreted as the ratio of the relative change in the halo radius to the relative change in the rate of lipid fragment production:

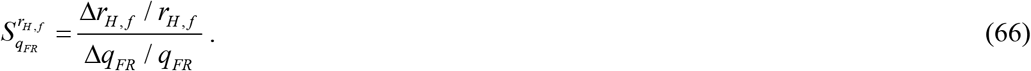

### 2.3. Numerical solution

Eqs. (6)-(8) and (36)-(38) were solved utilizing MATLAB’s ODE45 solver (MATLAB R2020b, MathWorks, Natick, MA, USA). To ensure the accuracy of the solution, the error tolerance parameters, RelTol and AbsTol, were set to 1e-10.

## 3. Results

Figs. 2-7 were generated with the assumption of zero initial concentrations of lipid membrane fragments and α-syn monomers. The data presented in Figs. 2-4 were calculated under the assumption of infinite half-lives for membrane fragments, fragment aggregates, α-syn monomers, and fibrils.

**Fig. 2.**
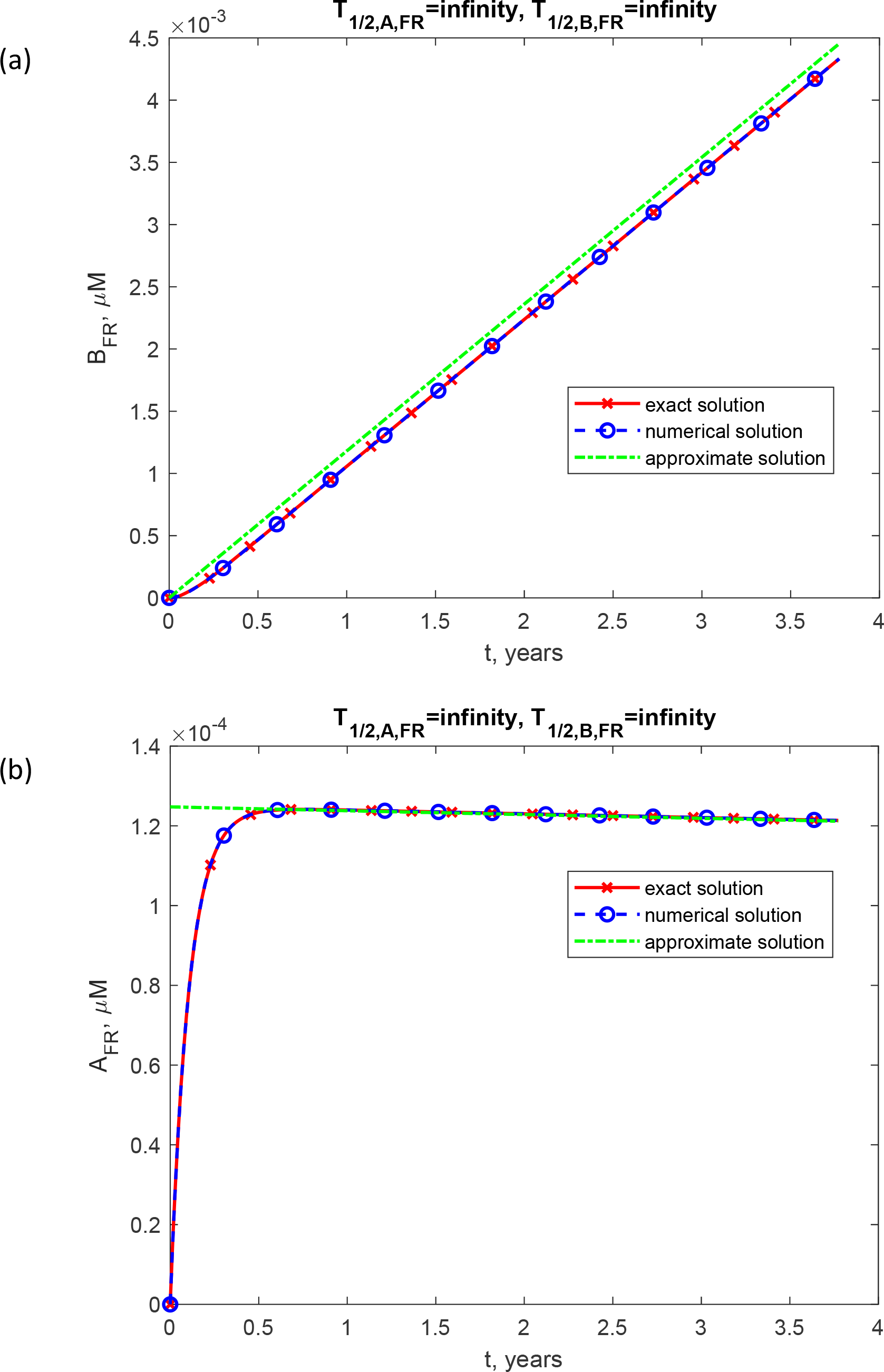
Molar concentration of lipid membrane aggregates, [*B*_*FR*_] (a) and lipid membrane fragments, [*A*_*FR*_] (b) as a function of time (in years). This scenario, depicted in the figure, assumes that both fragments and fragment aggregates have infinite half-lives. *q*_*FR*_ =1.57×10^−28^ mol s^-1^ and *q*_*AS*_ =1.47×10^−21^ mol s^-1^.

The concentration of lipid membrane aggregates exhibits a linear increase over time (proportional to both time, *t*, and the rate of membrane fragment production, *q*_*FR*_) as depicted in Fig. 2a and described by Eq. (16). On the other hand, the concentration of lipid membrane fragments initially experiences a rapid rise from zero but subsequently decreases over time as 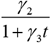, following the trend depicted in Fig. 2b and described by Eq. (19).

Fig. 2 also presents a comparison among numerical, exact, and approximate solutions for the concentrations of lipid membrane aggregates and lipid membrane fragments. As expected, the numerical and exact solutions agree perfectly. The approximate solution for the concentration of lipid membrane aggregates is only marginally different from the numerical solution. While the approximate solution strictly follows a linear pattern (as indicated by Eq. (16)), the numerical solution exhibits a subtle deviation from linearity, particularly during the initial phase of core growth, as shown in Fig. 2a. On the other hand, the numerical and approximate solutions for the concentration of lipid membrane fragments exhibit a disparity during the early stages of core growth, with the approximate solution violating the initial condition of zero concentration for the fragments. Nevertheless, these solutions nearly converge as time progresses, as depicted in Fig. 2b.

After the core of an LB is formed and the halo begins to grow, the concentration of α-syn fibrils demonstrates a linear increase over time, following the relationship 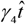, where 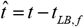 (Fig. 3a and Eq. (45)). The concentration of α-syn monomers initially exhibits a rapid increase from zero and subsequently decreases over time, following the relationship 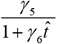 (Fig. 3b and Eq. (49)).

**Fig. 3.**
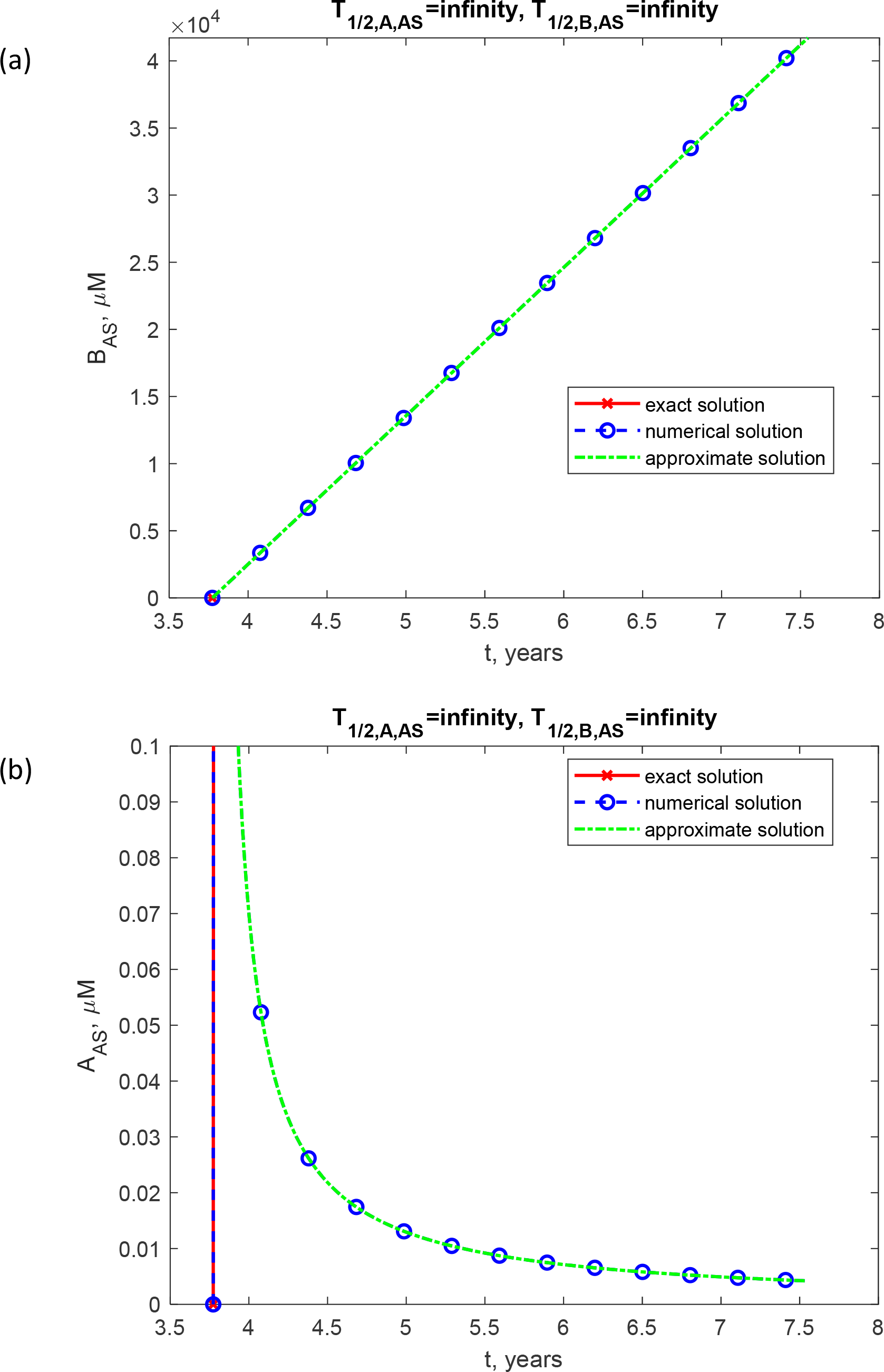
Molar concentration of α-syn fibrils, [*B*_*AS*_] (a) and α-syn monomers, [*A*_*AS*_] (b) as a function of time (in years). This scenario, depicted in the figure, assumes that both monomers and fibrils have infinite half-lives. *q*_*FR*_ =1.57×10^−28^ mol s^-1^ and *q*_*AS*_ =1.47×10^−21^ mol s^-1^.

In Fig. 3a and 3b, the numerical and approximate solutions almost coincide. An attempt to calculate an exact solution, given by Eq. (42), in Fig. 3, results in an NaN value. This occurs because the value of *q*_*AS*_ (1.47×10^−21^ mol s^-1^) used for Fig. 3 is seven orders of magnitude larger than the value of *q*_*FR*_ (1.57×10^−28^ mol s^-1^) used for Fig. 2. In order to verify the correctness of the exact solution in simulating the growth of the α-syn halo, computations reported in Fig. S1 were conducted using *q*_*AS*_ =1.47×10^−25^ mol s^-1^, which is four orders of magnitude smaller than the value in Fig. 3. Both the numerical and exact solutions in Fig. S1 show perfect agreement.

The LB core radius increases with time, following a cube root relationship, *a*_1_*t*^1/3^. After the LB core is formed and the halo starts growing, the halo radius increases with time, also following a cube root relationship, 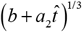. These trends are illustrated in Fig. 4 and described by Eqs. (31) and (61). The values of *q* (1.57×10^−28^ mol s^-1^) and *q* (1.47×10^−21^ mol s^-1^) are calculated to ensure the LB’s core and halo attain radii of 4 and 8 μm, respectively, which aligns with the observations in Fig. 4. These radii values are in line with the range of LB diameters reported in Olanow et al. (2004), which spans from 8 to 30 μm.

**Fig. 4.**
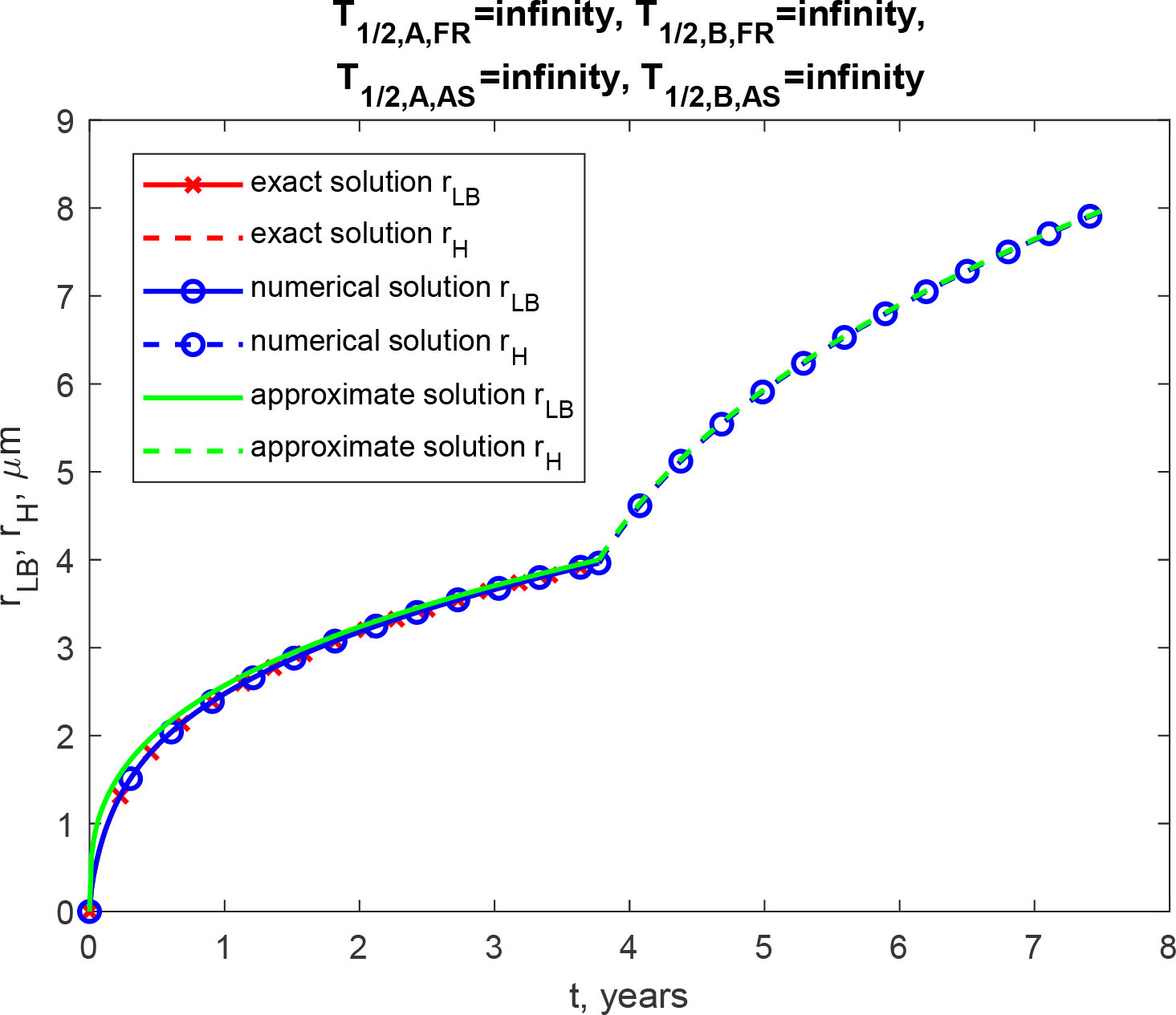
Radius of the growing core of the LB, *r*_*LB*_ (a) and the growing halo, *r*_*H*_ (b) over time (in years). This scenario, depicted in the figure, assumes that membrane fragments, fragment aggregates, α-syn monomers, and fibrils have infinite half-lives. *q*_*FR*_ =1.57×10^−28^ mol s^-1^ and *q*_*AS*_ =1.47×10^−21^ mol s^-1^.

In Fig. 4, the numerical and exact solutions closely match within the LB core region. However, in the halo region, attempting to compute the exact solution results in NAN in Eq. (42) due to the large value of *q*_*AS*_. The test of the exact solution is performed in Fig. S2, where the value of *q*_*AS*_ is reduced by four orders of magnitude. In Fig. S2, the numerical and exact solutions exhibit a perfect agreement in both the core and halo regions.

The data depicted in Figs. 5-7 were computed with the assumption of finite half-lives for membrane fragments, fragment aggregates, α-syn monomers, and fibrils, *T*_1/ 2, *A, FR*_ = 2.70 × 10^5^ s, *T*_1/ 2, *B, FR*_ = 1.35 ×10^6^ s, *T*_1/ 2, *A, AS*_ = 5.76 × 10^4^ s, and *T*_1/ 2, *B, AS*_ = 2.88 ×10^5^ s, respectively. The best effort was made to estimate physiologically relevant values of these parameters using information from existing literature. In Figs. 5-7, only numerical solutions are shown because the exact and approximate solutions are applicable exclusively to scenarios with infinite half-lives.

**Fig. 5.**
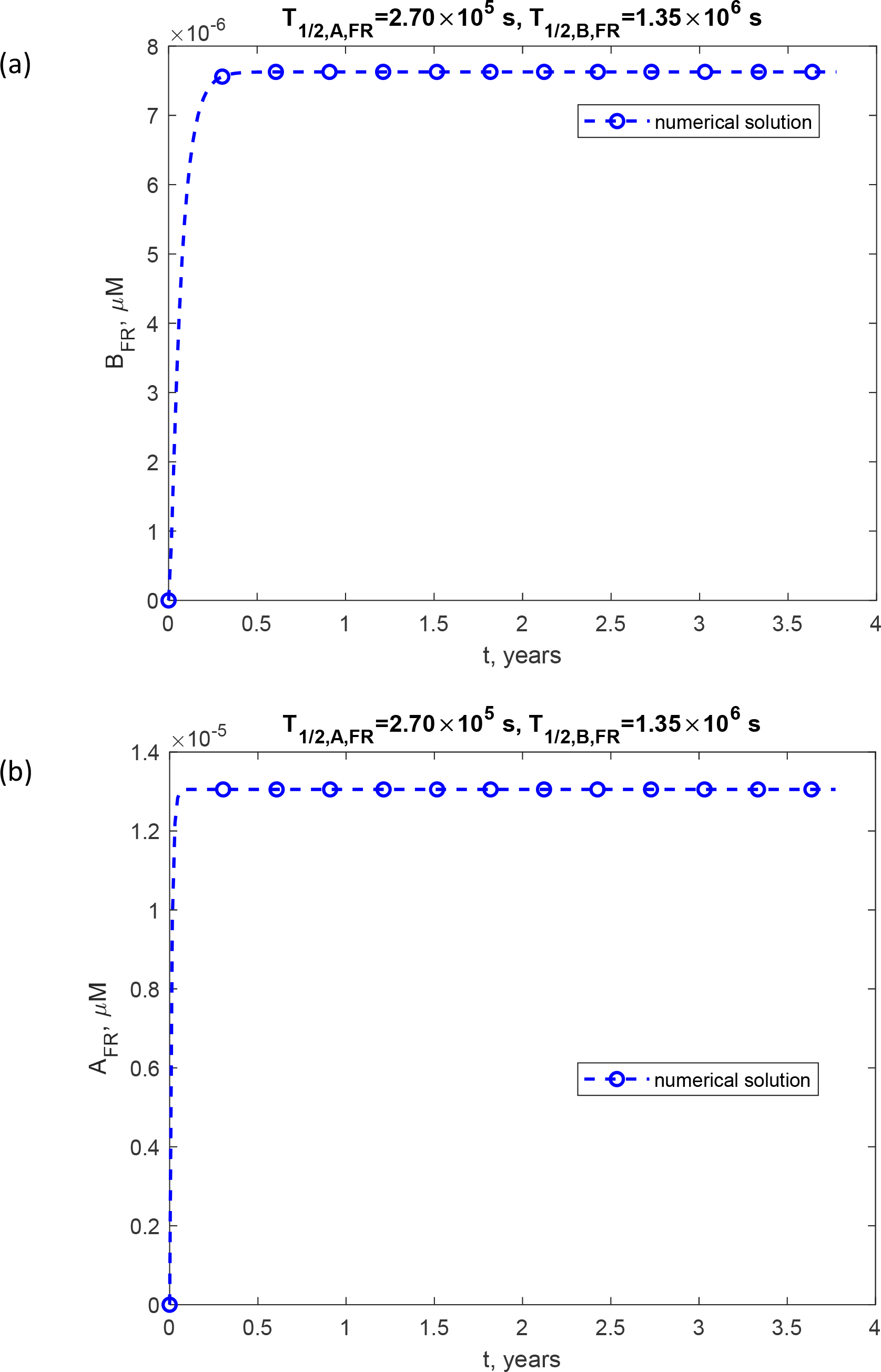
(a) Molar concentration of lipid membrane aggregates, [*B*_*FR*_] (a) and lipid membrane fragments, [*A*_*FR*_] (b) as a function of time (in years). This scenario, depicted in the figure, assumes that both fragments and fragment aggregates have finite half-lives, *T*_*1/2,A,FR*_ = 2.70 × 10^5^ s, *T*_*1/2,B,FR*_ = 1.35 ×10^6^ s, *q*_*FR*_ =1.57×10^−28^ mol s^-1^, and *q*_*AS*_ =1.47×10^−21^ mol s^-1^.

The concentrations of lipid membrane aggregates (Fig. 5a) and lipid membrane fragments (Fig. 5b) quickly approach their steady-state levels. This same behavior is observed for the concentrations of α-syn fibrils (Fig. 6a) and α-syn monomers (Fig. 6b). Initially, the core radius of the LB increases, primarily due to the initial rise in the concentration of lipid aggregates (as seen in Fig. 5a), and then stabilizes at a steady-state value. Similarly, once the core forms, the halo radius initially increases, primarily driven by the initial increase in the concentration of α-syn fibrils (as observed in Fig. 6a), before reaching a steady-state value (Fig. 7). The core radius measures approximately 0.5 μm, while the halo radius is approximately 1.15 μm. Comparing Fig. 4 (infinite half-lives of components and aggregates) with Fig. 7 (finite half-lives of components and aggregates), the LBs’ size in Fig. 4 is approximately 7 times larger than in Fig. 7.

**Fig. 6.**
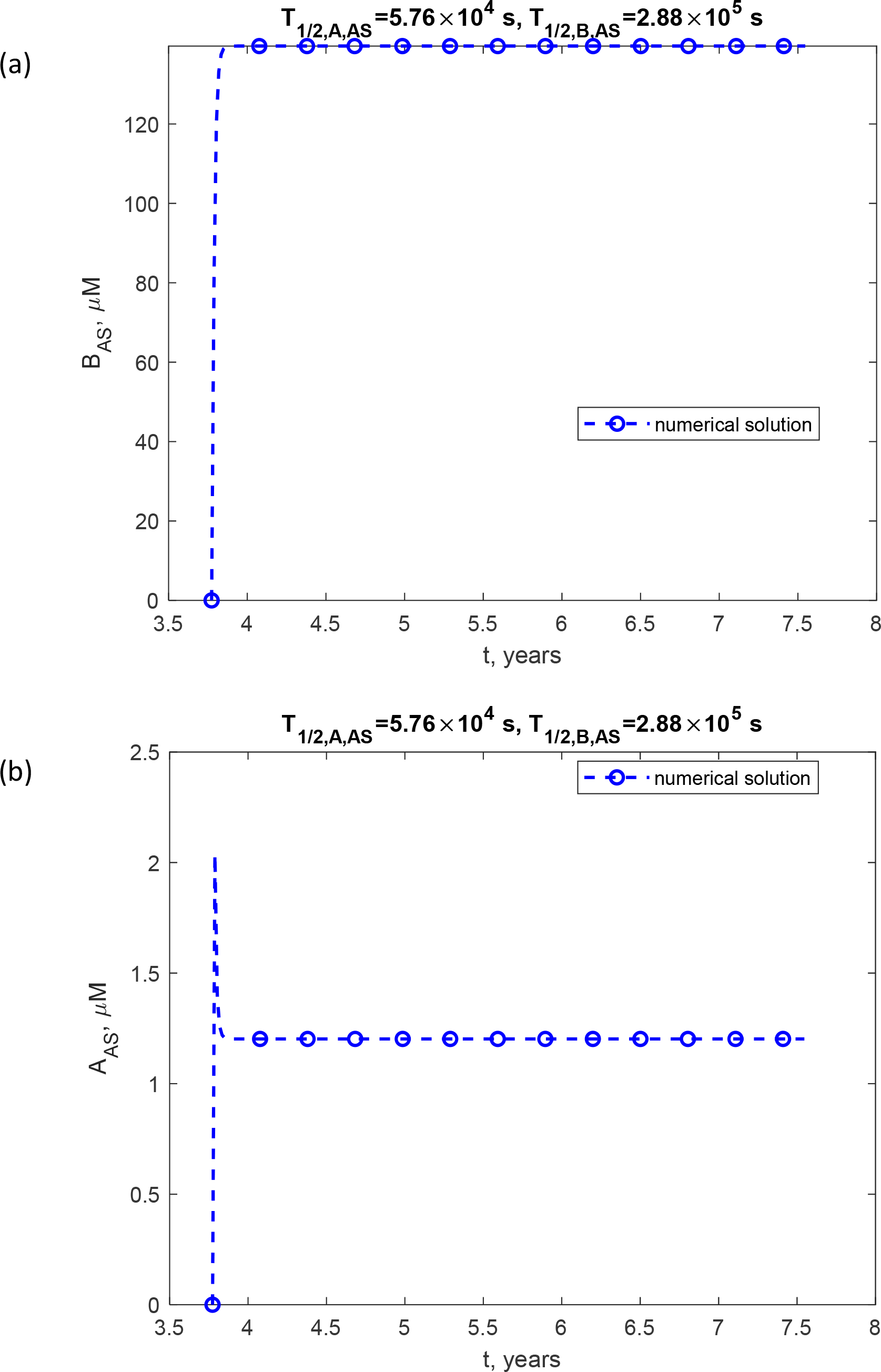
(a) Molar concentration of α-syn fibrils, [*B*_*AS*_] (a) and α-syn monomers, [*A*_*AS*_] (b) as a function of time (in years). This scenario, depicted in the figure, assumes that both monomers and fibrils have finite half-lives, *T*_*1/2,A,FR*_ = 5.76 × 10^4^ s, *T*_*1/2,B,FR*_ = 2.88 × 10^5^ s, *q*_*FR*_ =1.57×10^−28^ mol s^-1^, and *q*_*AS*_ =1.47×10^−21^ mol s^-1^.

**Fig. 7.**
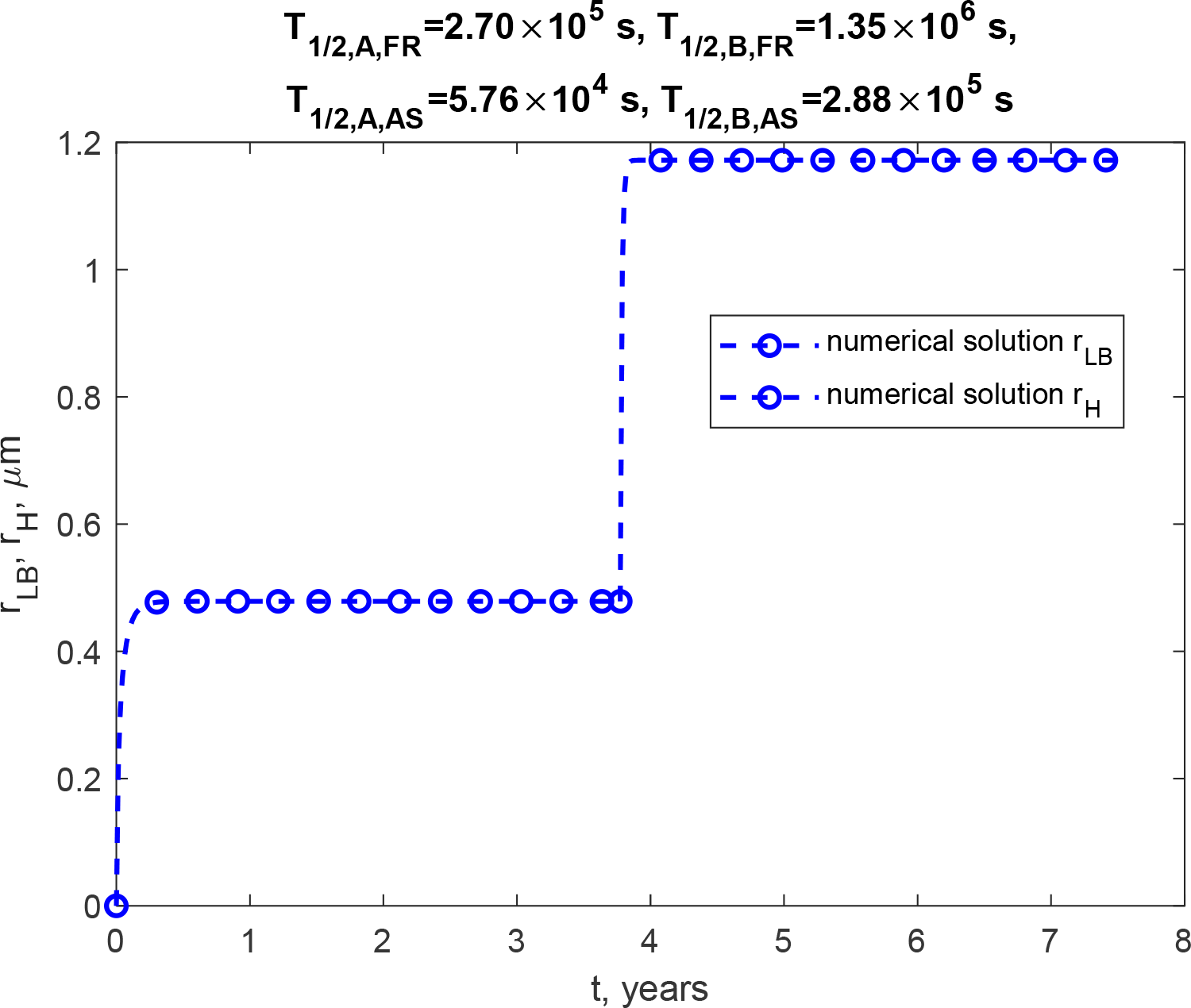
Radius of the growing core of the LB, *r*_*LB*_ (a) and the growing halo, *r*_*H*_ (b) over time (in years). This scenario, depicted in the figure, assumes that membrane fragments, fragment aggregates, α-syn monomers, and fibrils have finite half-lives, *T*_1/ 2, *A, FR*_ = 2.70 × 10^5^ s and *T*_1/ 2, *B, FR*_ = 1.35 × 10^6^ s, *T*_1/ 2, *A, AS*_ = 5.76 × 10^4^ s, *T*_1/ 2, *B, AS*_ = 2.88 × 10^5^ s, *q*_*FR*_ =1.57×10^−28^ mol s^-1^, and *q*_*AS*_ =1.47×10^−21^ mol s^-1^.

I asked whether the LB radius can be decreased to a size comparable to that displayed in Fig. 7 solely through the decay of membrane fragments and α-syn monomers, without accounting for the decay of lipid membrane aggregates and α-syn fibrils. This scenario corresponds to *T*_1/ 2, *B, FR*_ =∞ and *T*_1/ 2, *B, AS*_ =∞. The scenario with *T*_1/ 2, *A, FR*_ = 2.70 × 10^5^ s and *T*_1/ 2, *A, AS*_ = 5.76 ×10^4^ s (the same values as those used for Fig. 7) is presented in Figs. S3-S5 in the Supplemental Materials. In this scenario, the halo radius after 7.5 years reaches approximately 5.5 μm (Fig. S3), which is just 0.5 μm less than that in Fig. 7. However, if *T*_1/ 2, *A, FR*_ = 2.70 × 10^2^ s and *T*_1/ 2, *A, AS*_ = 5.76 ×10^1^ s (the values decreased by a factor of 1000) are employed, the halo radius after 7.5 years reaches approximately 1.15 μm, the same value as that in Fig. 7. Thus, high decay rates of membrane fragments and α-syn monomers can slow down the growth of LBs, without the decay of lipid membrane aggregates and α-syn fibrils.

The sensitivity of the fully grown LB core radius, *r*_*LB, f*_, to the rate of membrane fragment production, *q*_*FR*_, is positive (Tables 6 and 7). This indicates that when fragments are generated at a higher rate, the LB core tends to grow to a larger size. The value of 0.334 for 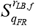 can be understood by using the approximate solution given by Eq. (29):

**Table 6.**
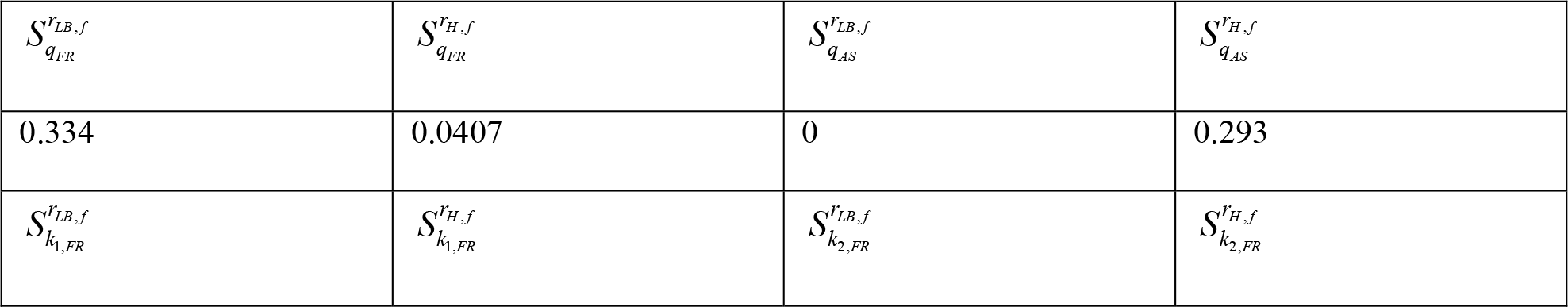
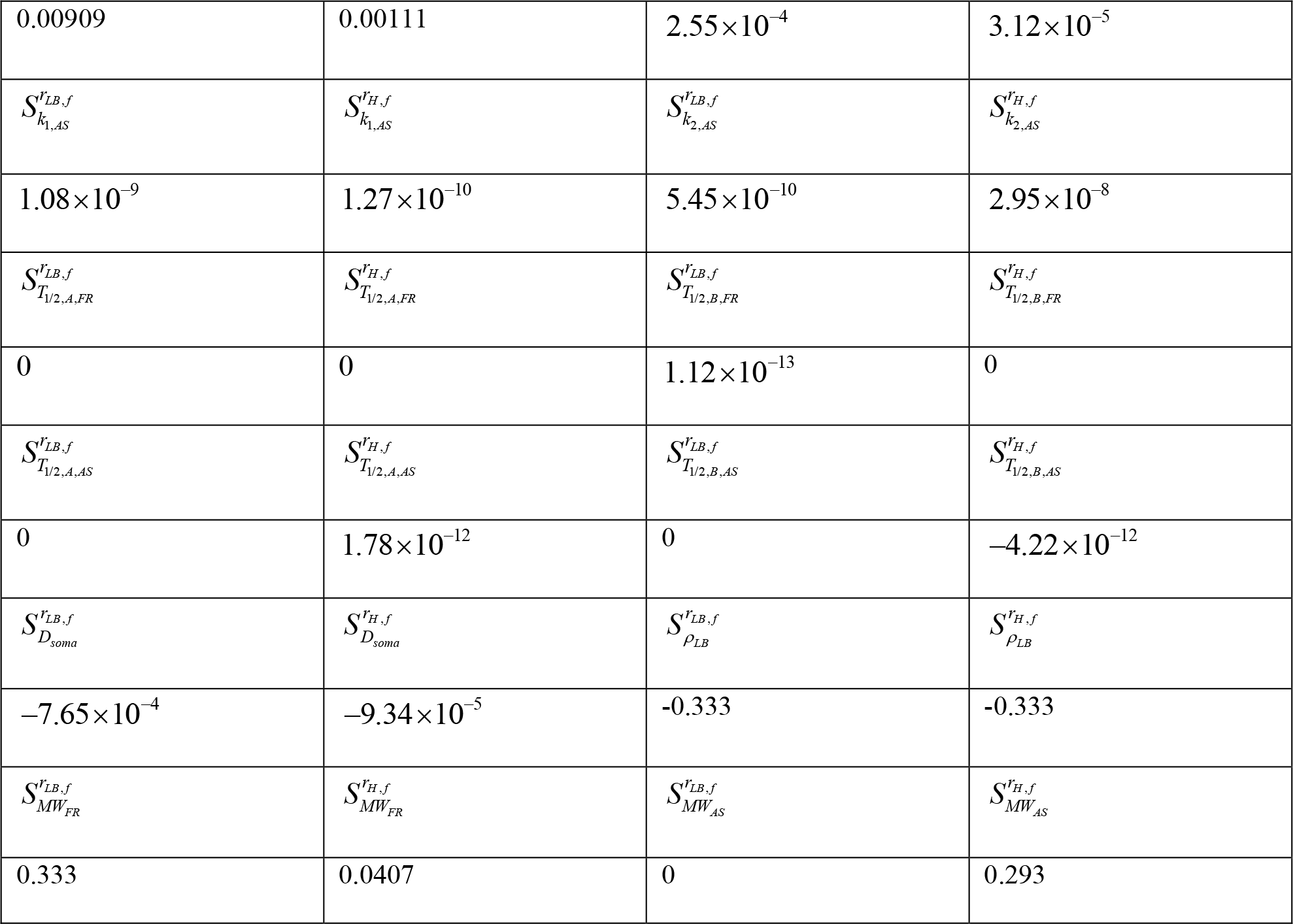
Relative sensitivity of the radii of the fully grown LB core, *r*_*LB, f*_, and the fully grown LB halo, *r*_*H, f*_, to various model parameters. Computations were carried out using Eqs. (64) and (65) with (for example) Δ*q*_*FR*_ = 10^−3^ *q*_*FR*_. The table is computed for the scenario where membrane fragments, fragment aggregates, α-syn monomers, and fibrils have infinite half-lives, *T*_1/ 2, *A, FR*_ →∞, *T*_1/ 2, *B, FR*_ →∞, *T*_1/ 2, *A, AS*_ →∞, and *T*_1/ 2, *B, AS*_ →∞.

**Table 7.**
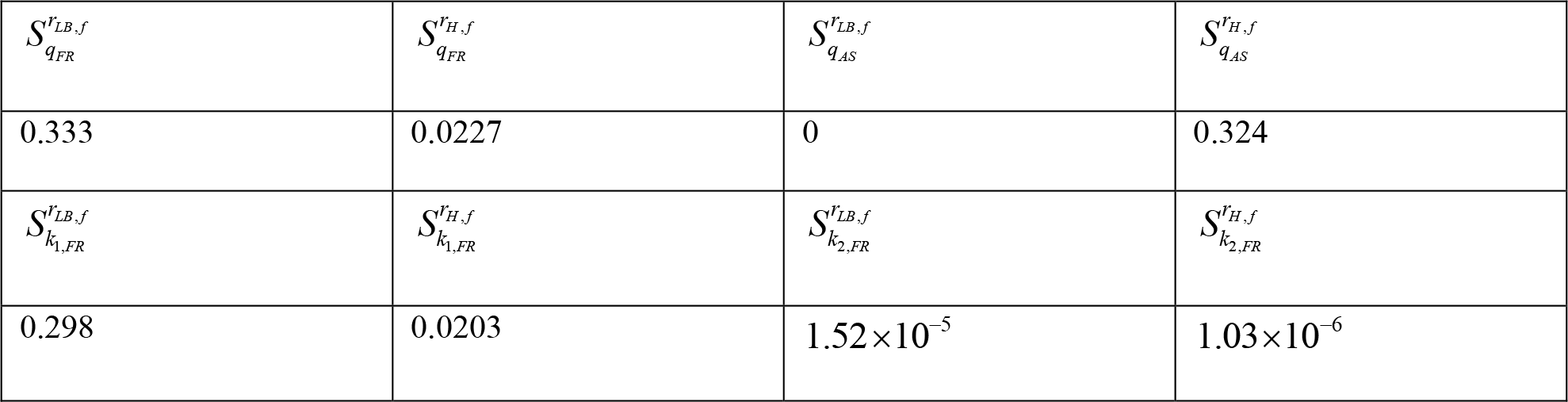
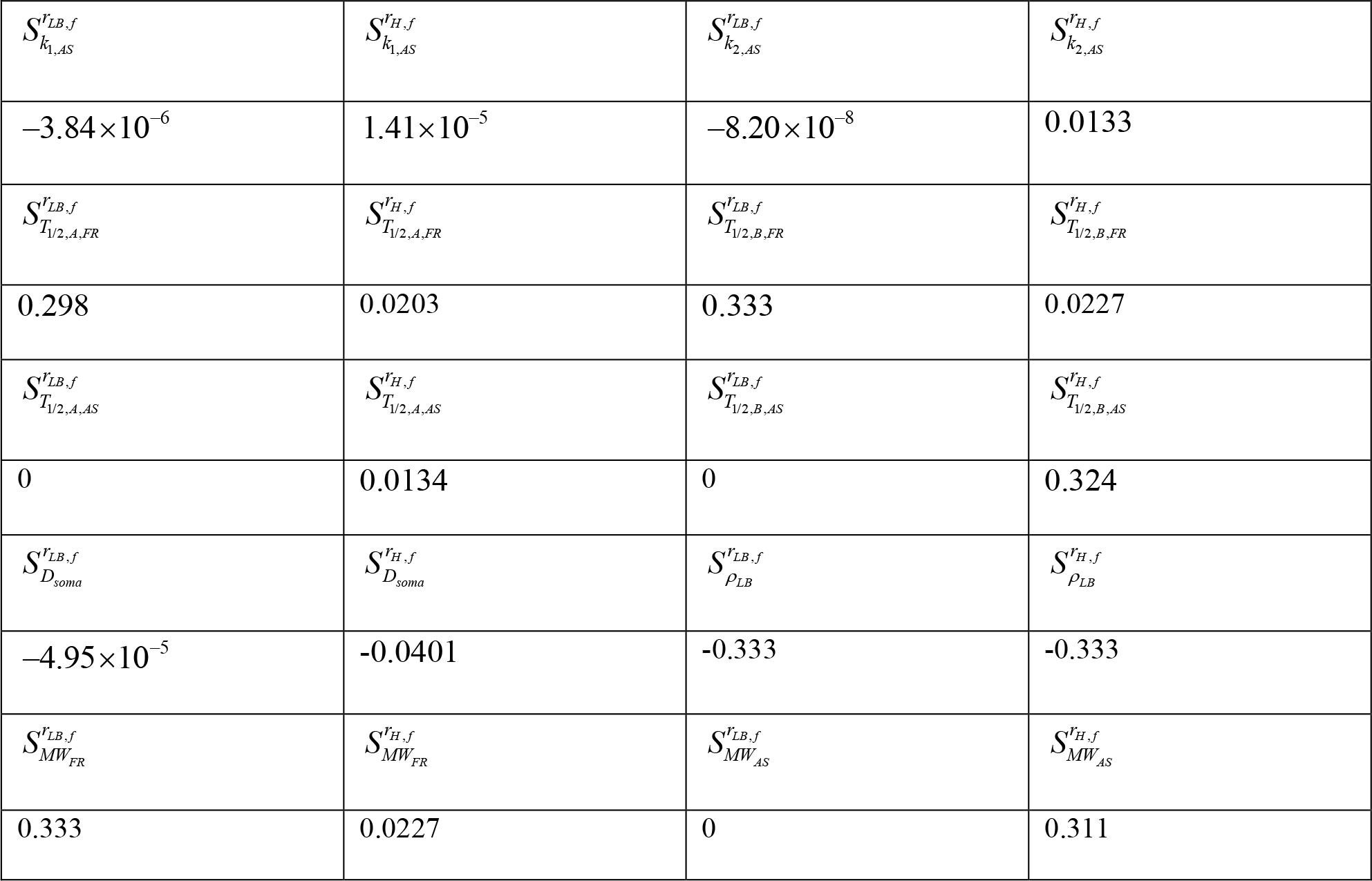
Relative sensitivity of the radii of the fully grown LB core, *r*_*LB, f*_, and the fully grown LB halo, *r*_*H, f*_, to various model parameters. Computations were carried out using Eqs. (64) and (65) with (for example) Δ*q*_*FR*_ = 10^−3^ *q*_*FR*_. The table is computed for the scenario where membrane fragments, fragment aggregates, α-syn monomers, and fibrils have finite half-lives, *T*_1/ 2, *A, FR*_ = 2.70 × 10^5^ s, *T*_1/ 2, *B, FR*_ = 1.35 ×10^6^ s, *T*_1/ 2, *A, AS*_ = 5.76 × 10^4^ s, and *T*_1/ 2, *B, AS*_ = 2.88 × 10^5^ s.

|  |  |  |  |
| --- | --- | --- | --- |
| $S_{q_{FR}}^{r_{LB},f}$ | $S_{q_{FR}}^{r_{H},f}$ | $S_{q_{AS}}^{r_{LB},f}$ | $S_{q_{AS}}^{r_{H},f}$ |
| 0.333 | 0.0227 | 0 | 0.324 |
| $S_{k_{1,FR}}^{r_{LB},f}$ | $S_{k_{1,FR}}^{r_{H},f}$ | $S_{k_{2,FR}}^{r_{LB},f}$ | $S_{k_{2,FR}}^{r_{H},f}$ |
| 0.298 | 0.0203 | $1.52 \times 10^{-5}$ | $1.03 \times 10^{-6}$ |
| $S_{k_{1,AS}}^{r_{LB},f}$ | $S_{k_{1,AS}}^{r_{H},f}$ | $S_{k_{2,AS}}^{r_{LB},f}$ | $S_{k_{2,AS}}^{r_{H},f}$ |
| $-3.84 \times 10^{-6}$ | $1.41 \times 10^{-5}$ | $-8.20 \times 10^{-8}$ | 0.0133 |
| $S_{T_{1/2,A,FR}}^{r_{LB},f}$ | $S_{T_{1/2,A,FR}}^{r_{H},f}$ | $S_{T_{1/2,B,FR}}^{r_{LB},f}$ | $S_{T_{1/2,B,FR}}^{r_{H},f}$ |
| 0.298 | 0.0203 | 0.333 | 0.0227 |
| $S_{T_{1/2,A,AS}}^{r_{LB},f}$ | $S_{T_{1/2,A,AS}}^{r_{H},f}$ | $S_{T_{1/2,B,AS}}^{r_{LB},f}$ | $S_{T_{1/2,B,AS}}^{r_{H},f}$ |
| 0 | 0.0134 | 0 | 0.324 |
| $S_{D_{soma}}^{r_{LB},f}$ | $S_{D_{soma}}^{r_{H},f}$ | $S_{\rho_{LB}}^{r_{LB},f}$ | $S_{\rho_{LB}}^{r_{H},f}$ |
| $-4.95 \times 10^{-5}$ | -0.0401 | -0.333 | -0.333 |
| $S_{MW_{FR}}^{r_{LB},f}$ | $S_{MW_{FR}}^{r_{H},f}$ | $S_{MW_{AS}}^{r_{LB},f}$ | $S_{MW_{AS}}^{r_{H},f}$ |
| 0.333 | 0.0227 | 0 | 0.311 |

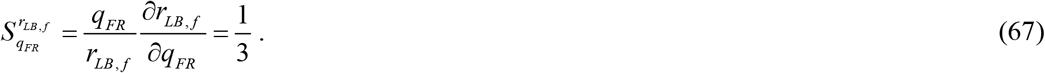

The positive sensitivity of the halo’s radius, *r*_*H, f*_, to *q*_*FR*_ is a result of the halo forming on top of the core (as shown in Fig. 1). Therefore, when the core is larger, it leads to a larger halo. The positive sensitivity of the halo’s radius, *r*_*H, f*_, to the rate of α-syn monomer production, *q*_*AS*_, is due to the fact that larger *q*_*AS*_ results in a faster material supply for the halo buildup. *r*_*LB, f*_ is unaffected by *q*_*AS*_ because α-syn monomers do not play a role in the formation of the LB core. The observed trends hold true for both scenarios, whether considering infinite (Table 6) or finite (Table 7) half-lives of components and aggregates.

For scenarios with infinite half-lives of components and aggregates (Table 6), the sensitivities of *r*_*LB, f*_ and *r*_*H, f*_ to the kinetic constants, *k*_1,*FR*_, *k*_2,*FR*_, *k*_1, *AS*_, and *k*_2, *AS*_ are nearly zero. This is consistent with the approximate solutions for the core and halo radii given by Eqs. (29) and (59), respectively, which indicate an absence of dependence on the kinetic constants. In the case of finite half-lives for components and aggregates, the approximate solutions are not applicable. The core and halo radii, *r*_*LB, f*_ and *r*_*H, f*_, exhibit greater sensitivity to the kinetic constants, especially to *k*_1, *FR*_ (Table 7).

When considering infinite half-lives for components and aggregates, both *r*_*LB, f*_ and *r*_*H, f*_ show no sensitivity to changes in the half-lives, *T*_1/ 2, *A,FR*_, *T*_1/ 2,*B,FR*_, *T*_1/ 2, *A, AS*_, and *T*_1/ 2,*B, AS*_ (since an infinitely large value remains infinitely large even when varied by 0.1%). With finite half-lives for components and aggregates, both *r*_*LB, f*_ and *r*_*H, f*_ demonstrate a considerably higher sensitivity to the half-lives, particularly to *T*_1/ 2, *A,FR*_ and *T*_1/ 2,*B,FR*_ (Table 7). This is due to the fact that when the half-lives are finite, the core and halo radii quickly attain their steady-state values (as depicted in Fig. 7). These values are determined by the equilibrium between the rates of membrane fragments and α-syn monomers production and their respective decay rates. The sensitivities to *T*_1/ 2, *A,FR*_ and *T*_1/ 2,*B,FR*_ are positive (Table 7) because an increase in the half-life results in a reduced decay rate of lipid membrane fragments and fragment aggregates, as evident from the last terms on the right-hand side of Eqs. (3) and (4). Because α-syn monomers do not play a role in forming the LB core, the sensitivity of *r*_*LB, f*_ to *T*_1/ 2, *A, AS*_ and *T*_1/ 2,*B, AS*_ is zero (as shown in Table 7).

In the case of infinite half-lives of components and aggregates (Table 6), both *r*_*LB, f*_ and *r*_*H, f*_ remain unaffected by changes in the soma diameter, *D*_*soma*_ (within the limits of the approximate solution). This agrees with the approximate solutions given by Eqs. (29) and (59), which indicate that neither *r*_*LB, f*_ nor *r*_*H, f*_ are affected by variations in *D*_*soma*_. However, when considering finite half-lives for components and aggregates (Table 7), *r*_*H, f*_ does exhibit a slight negative sensitivity to changes in *D*_*soma*_.

The sensitivity of both *r*_*LB, f*_ and *r*_*H, f*_ to the density of the LB, *ρ*_*LB*_, is found to be -0.333. To understand these relationships, let us refer to Eq. (29):

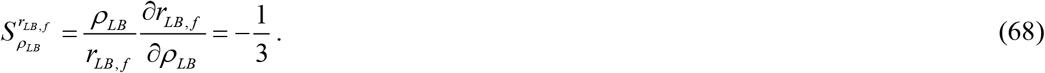

Utilizing Eq. (59):

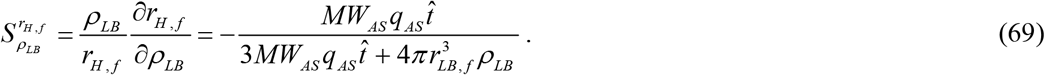

In the limit of 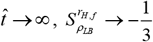.

The sensitivity of the core radius, *r*_*LB, f*_, to the parameter *MW*_*FR*_, is found to be 0.333, as shown in Tables 6 and 7. This relationship can be explained by employing the approximate solution provided in Eq. (29):

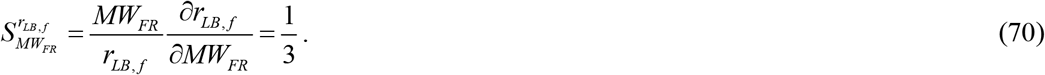

The sensitivity of the halo radius, *r*_*H, f*_, to *MW*_*FR*_ is slightly positive (Tables 6 and 7). In contrast, the sensitivity of *r*_*H, f*_ to *MW*_*FR*_ is zero, which happens because α-syn does not affect the formation of the LB core. Additionally, the sensitivity of *r*_*H, f*_ to *MW*_*AS*_ is positive (Tables 6 and 7).

## 4. Discussion, limitations of the model, and future directions

The primary finding of this study is the “cube root hypothesis.” According to this hypothesis, the core size of the LB, *r*_*LB,f*_, increases in a manner proportional to the cube root of time, *a*_1_*t*^1/3^. Additionally, the radius of LB halo, *r*_*H, f*_, which starts growing after the core formation is complete (this moment corresponds to 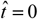), increases in a manner proportional to 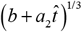. The cube root hypothesis is only applicable when the degradation machinery for lipid fragments and α-syn fails, resulting in infinitely long half-lives for these components. It is worth stressing that, even though the mathematical analysis leading to the cube root result (as demonstrated in the derivation of Eqs. (29) and (59)) is complex, the fundamental physics underlying this result is remarkably elegant. The rationale behind the cube root relationship is that, in the absence of the decay of membrane fragment and α-syn components and aggregates, LB growth is primarily governed by the rates at which lipid fragments and α-syn monomers are produced. Assuming these rates remain constant, the increase in LB volume becomes linear with time, consequently leading to the LB core radius growing in proportion to the cube root of time.

The approximate solutions made it possible to determine the production rates of lipid membrane fragments, denoted as *q*_*FR*_, and α-syn monomers, denoted as *q*_*AS*_, necessary to form an LB with specified core and halo radii, *r*_*LB, f*_ and *r*_*H, f*_, respectively. For the scenario when the degradation mechanisms for lipids and α-syn malfunction (resulting in infinitely long half-lives for lipid and α-syn components and aggregates), computations were conducted to model the growth of an LB that achieves a core radius of 4 μm and a halo radius of 8 μm within a 7.5-year span of growth. These values align with previously reported LB diameters (ranging from 8 to 30 μm, as reported in Olanow et al. (2004)).

When the lipid and α-syn degradation processes are operational, with half-lives reflecting physiological values, over a 7.5-year span of growth the core radius reaches approximately 0.5 μm and the halo radius reaches approximately 1.15 μm. Consequently, the predicted size of the LBs in this case is approximately 7 times smaller than that predicted for the scenario with dysfunctional degradation machinery.

The approximate solutions for the LB core and halo radii, derived under the assumption of dysfunctional degradation machinery (infinite half-lives), were validated through comparison with numerical and exact analytical solutions. These approximate solutions for the LB core and halo radii can be used for fitting future experimental data.

The sensitivity analysis based on numerical solutions confirmed that in scenarios with infinite half-lives of components and aggregates, the radii of the LB core and halo do not depend on the kinetic constants that describe the rates of nucleation and autocatalytic growth of lipid fragment aggregates and α-syn fibrils. This conclusion agrees with the findings obtained using the approximate solutions.

The obtained results have limitations because they rely on the simplified F-W aggregation model, which does not differentiate between aggregates of different sizes. Given that LBs are composed of more than 300 proteins according to Shahmoradian et al. (2019), there is a need for more sophisticated models that account for the intricate chain of aggregation processes involved in LB formation.

Future studies should focus on mechanistic modeling of the production and transport of lipid membrane fragments and α-syn monomers. Additionally, it is crucial to explore the connection between α-syn overexpression and other pathological processes within neurons, which may contribute to the formation of LBs. In particular, an excess of α-syn and especially α-syn oligomers have been observed to trigger mitochondrial fragmentation (Stefanis, 2012; Krzystek et al., 2021). This is relevant to LB formation since fragmented and distorted mitochondria have been found in LBs, as noted in Shahmoradian et al. (2019). Future models should consider the potential spread of LB pathology (Fares et al., 2021).

In future research, it is also important to create a physical model that elucidates the factors behind the cessation of lipid core growth and the initiation of halo formation by radiating filaments on its surface. This phenomenon is likely attributable to thermodynamic instability. An analysis of this instability should enable the prediction of the critical core radius at which the halo begins to develop on the core’s surface. Future models should take into consideration the influence of LB size on the rate of LB surface growth (Martin, 2020).

## Abbreviations

Aβ: amyloid-β
α-syn: alpha-synuclein
F-W: Finke-Watzky
LB: Lewy body
PD: Parkinson’s disease
SN: substantia nigra

## Acknowledgment

AVK acknowledges the support provided by the National Science Foundation (grant CBET-2042834) and the Alexander von Humboldt Foundation through the Humboldt Research Award.

## Supplemental Materials

### S1. Supplemental figures

**Fig. S1.**
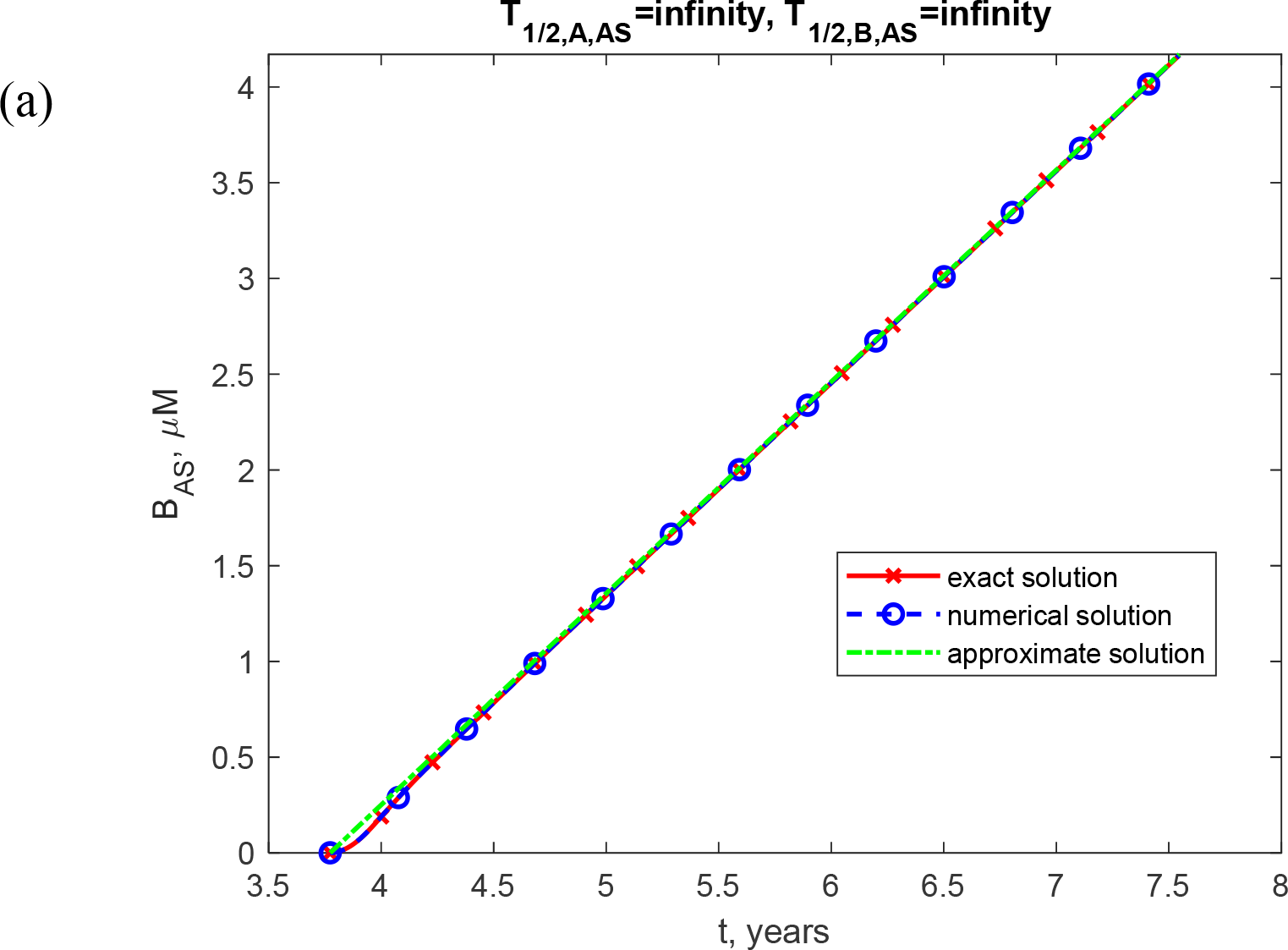
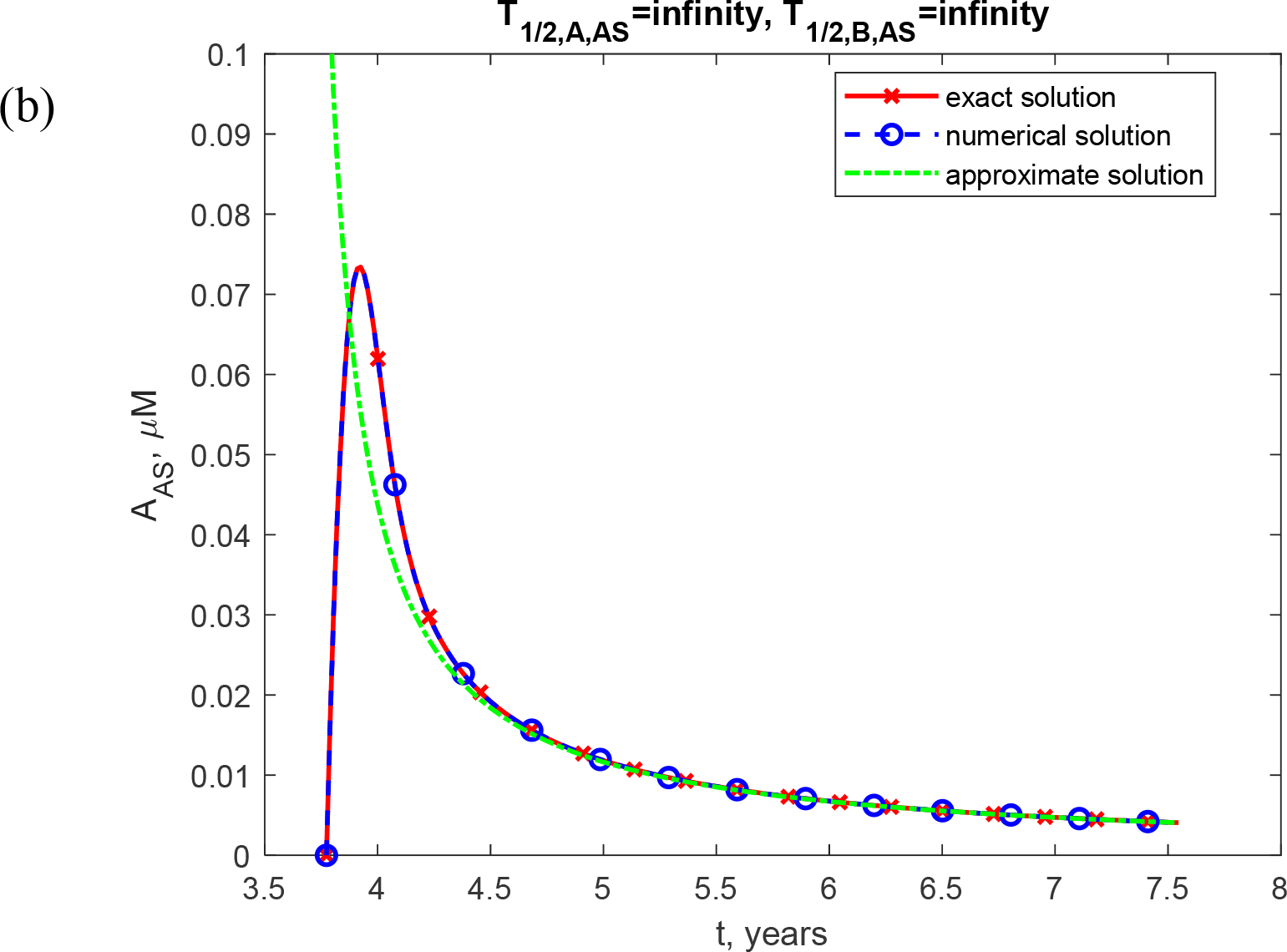
Molar concentration of α-syn fibrils, [*B*_*AS*_] (a) and α-syn monomers, [*A*_*AS*_] (b) as a function of time (in years). This scenario, depicted in the figure, assumes that both monomers and fibrils have infinite half-lives. *q*_*FR*_ =1.57×10^−28^ mol s^-1^ and *q*_*AS*_ =1.47×10^−25^ mol s^-1^ (*q*_*AS*_ is four orders of magnitude smaller than the value used for calculating Figs. 1-7).

**Fig. S2.**
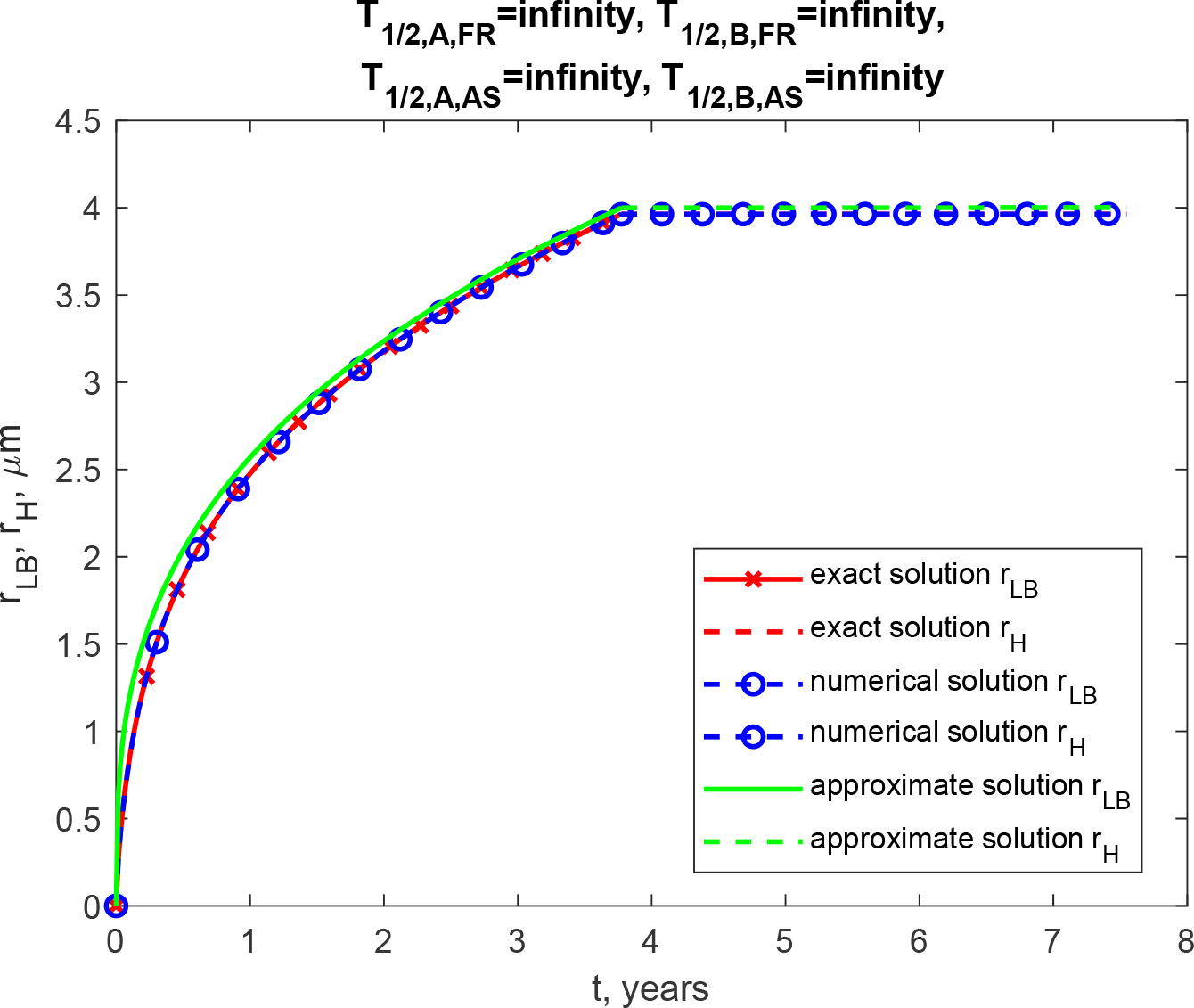
Radius of the growing core of the LB, *r*_*LB*_ (a) and the growing halo, *r*_*H*_ (b) over time (in years). This scenario, depicted in the figure, assumes that membrane fragments, fragment aggregates, α-syn monomers, and fibrils have infinite half-lives. *q*_*FR*_ =1.57×10^−28^ mol s^-1^ and *q*_*AS*_ =1.47×10^−25^ mol s^-1^ (*q*_*AS*_ is four orders of magnitude smaller than the value used for calculating Figs. 1-7).

**Fig. S3.**
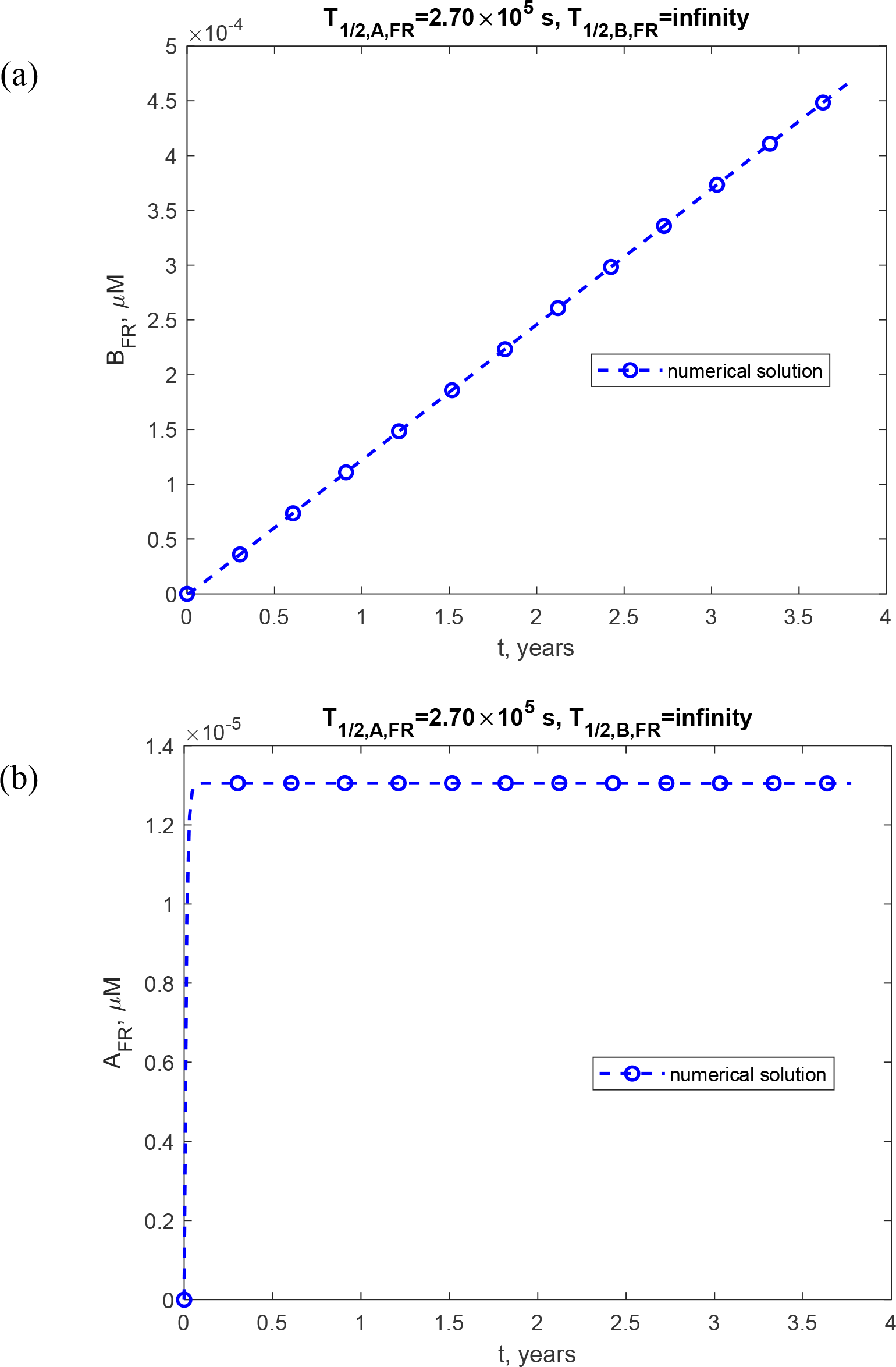
(a) Molar concentration of lipid membrane aggregates, [*B*_*FR*_] (a) and lipid membrane fragments, [*A*_*FR*_] (b) as a function of time (in years). This scenario, depicted in the figure, assumes that fragments have a finite half-life, *T*_1/ 2, *A, FR*_ = 2.70 × 10^5^ s, while fragment aggregates have an infinite half-life. *q*_*FR*_ =1.57×10^−28^ mol s^-1^ and *q*_*AS*_ =1.47×10^−21^ mol s^-1^.

**Fig. S4.**
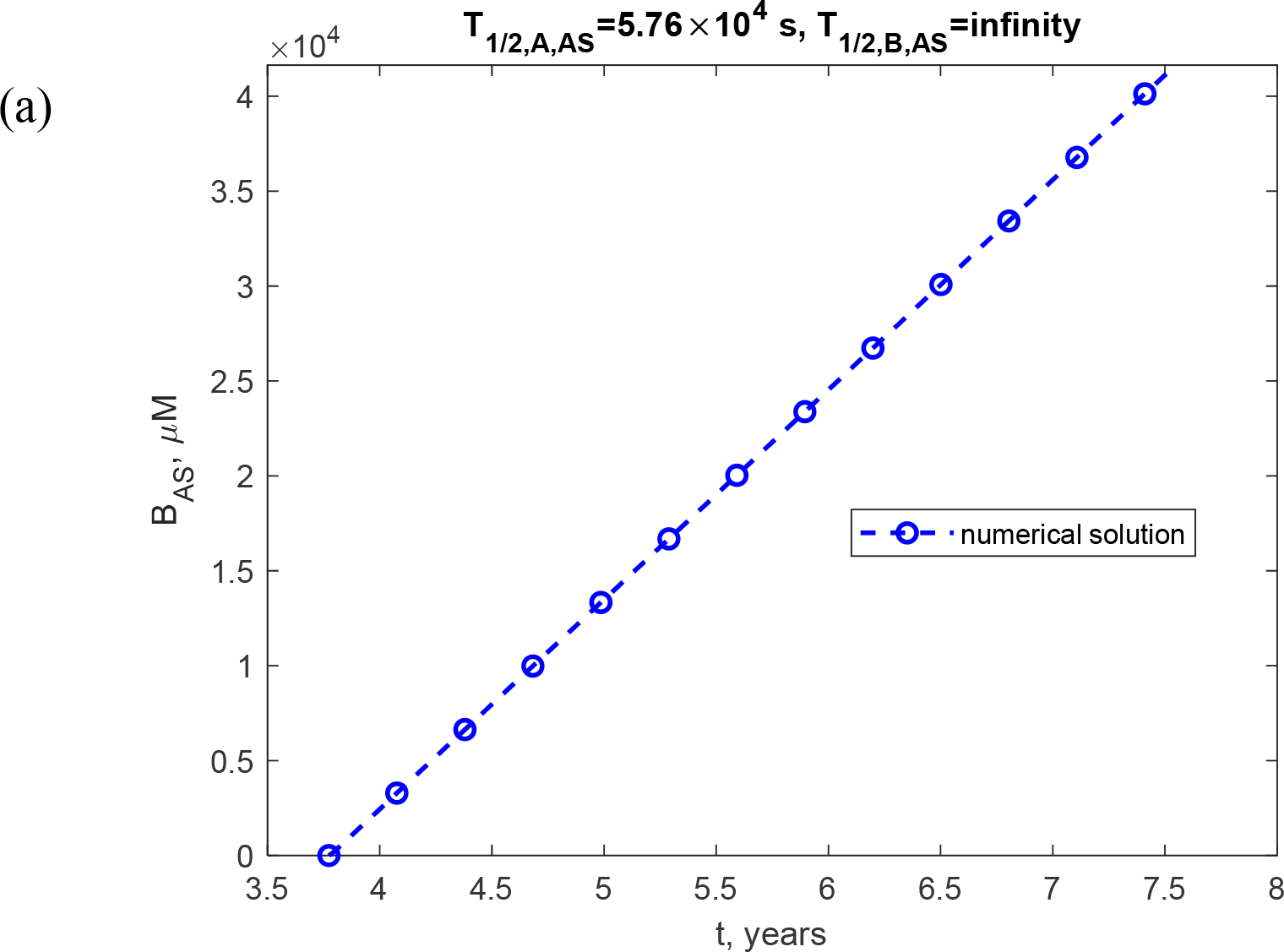
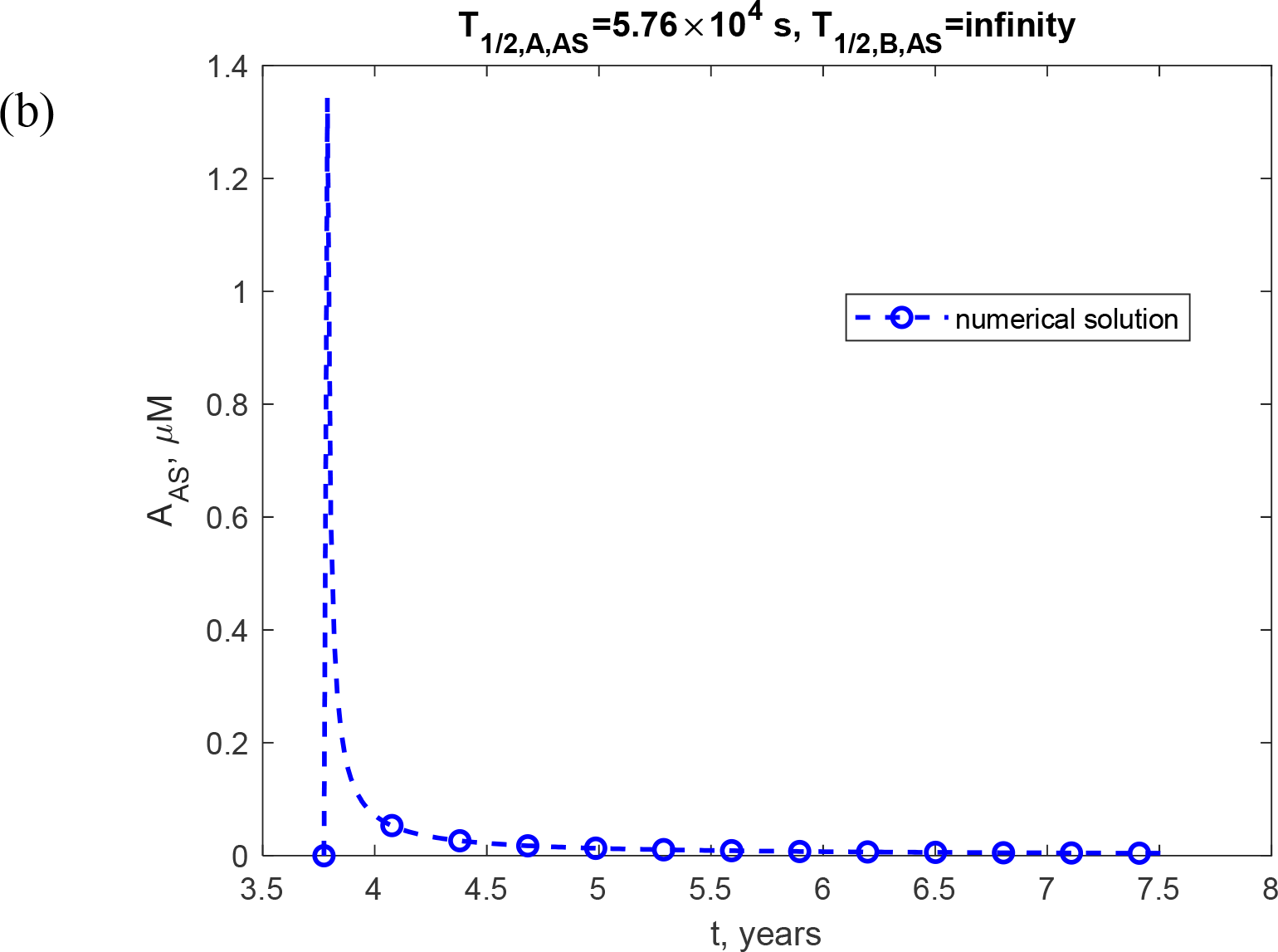
(a) Molar concentration of α-syn fibrils, [*B*_*AS*_] (a) and α-syn monomers, [*A*_*AS*_] (b) as a function of time (in years). This scenario, depicted in the figure, assumes that α-syn monomers have a finite half-life, *T*_1/ 2, *A, AS*_ = 5.76 × 10^4^ s, while α-syn fibrils have an infinite half-life. *q*_*FR*_ =1.57×10^−28^ mol s^-1^ and *q*_*AS*_ =1.47×10^−21^ mol s^-1^.

**Fig. S5.**
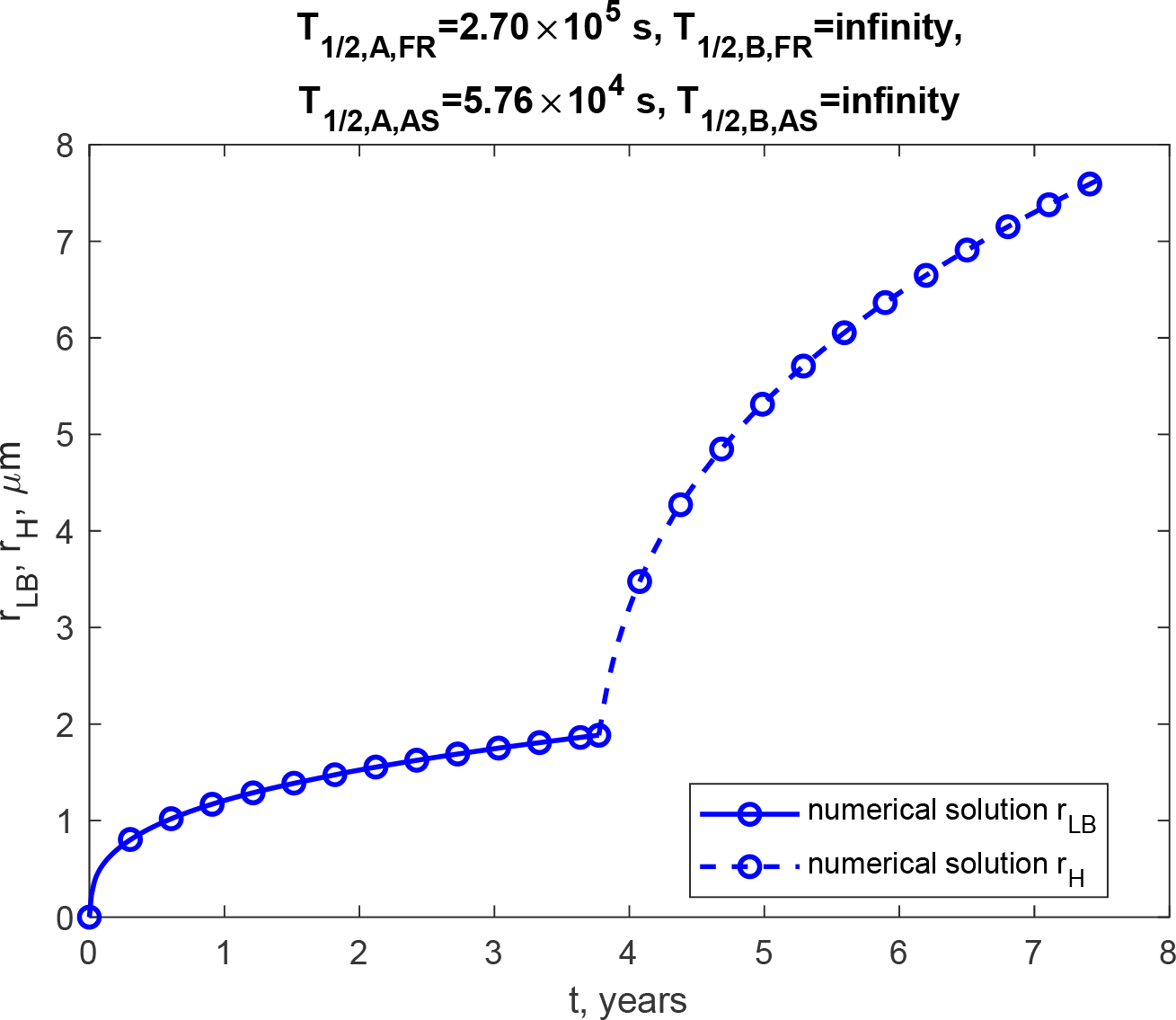
Radius of the growing core of the LB, *r*_*LB*_ (a) and the growing halo, *r*_*H*_ (b) over time (in years). This scenario, depicted in the figure, assumes that membrane fragments and α-syn monomers have finite half-life, *T*_1/ 2, *A, FR*_ = 2.70 × 10^5^ s and *T*_1/ 2, *A, AS*_ = 5.76 ×10^4^ s, while fragment aggregates and α-syn fibrils have infinite half-lives. *q*_*FR*_ =1.57×10^−28^ mol s^-1^ and *q*_*AS*_ =1.47×10^−21^ mol s^-1^.

**Fig. S6.**
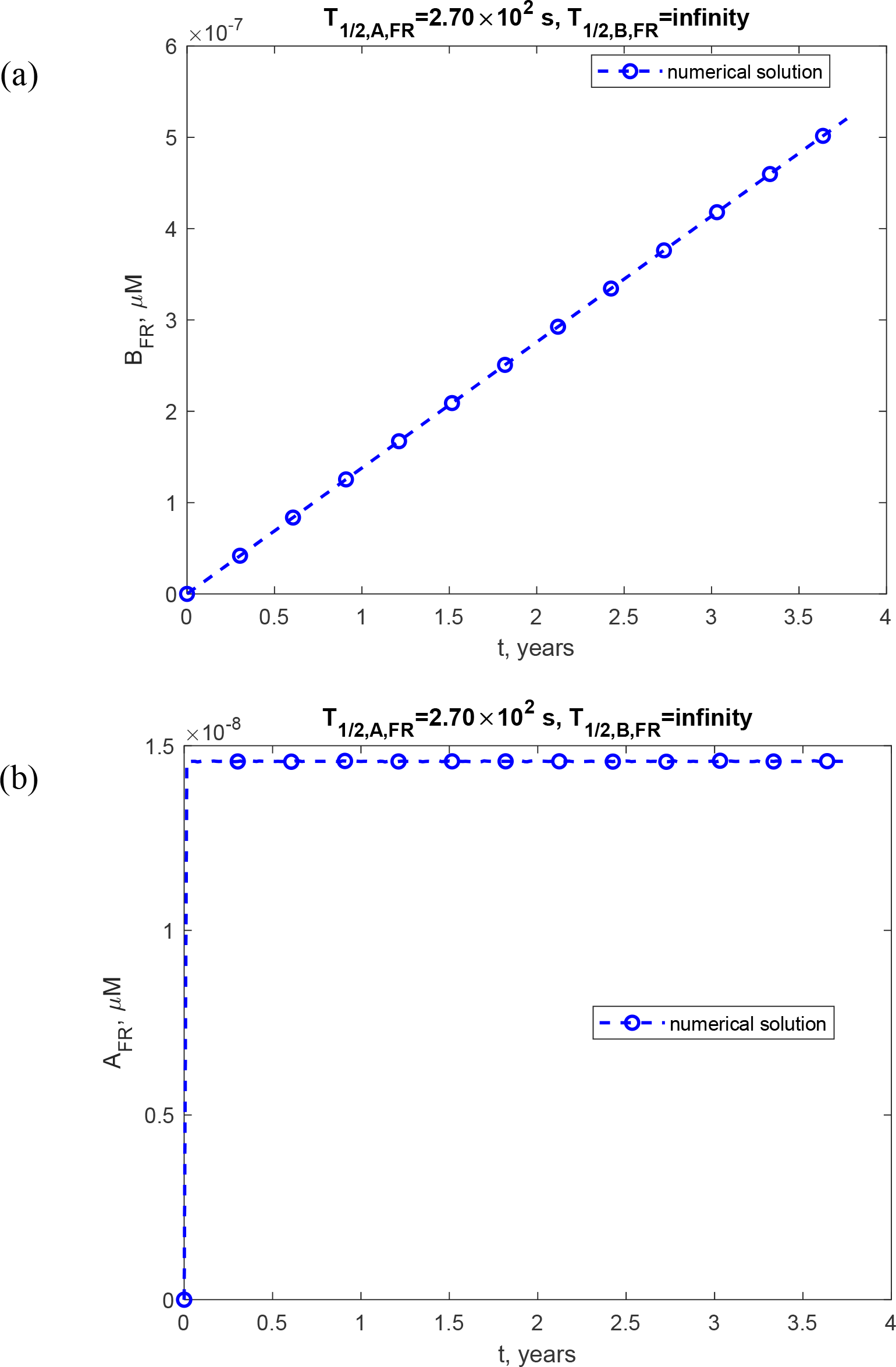
(a) Molar concentration of lipid membrane aggregates, [*B*_*FR*_] (a) and lipid membrane fragments, [*A*_*FR*_] (b) as a function of time (in years). This scenario, depicted in the figure, assumes that fragments have a finite half-life, *T*_1/ 2, *A, FR*_ = 2.70 × 10^2^ s, while fragment aggregates have an infinite half-life. *q*_*FR*_ =1.57×10^−28^ mol s^-1^ and *q*_*AS*_ =1.47×10^−21^ mol s^-1^.

**Fig. S7.**
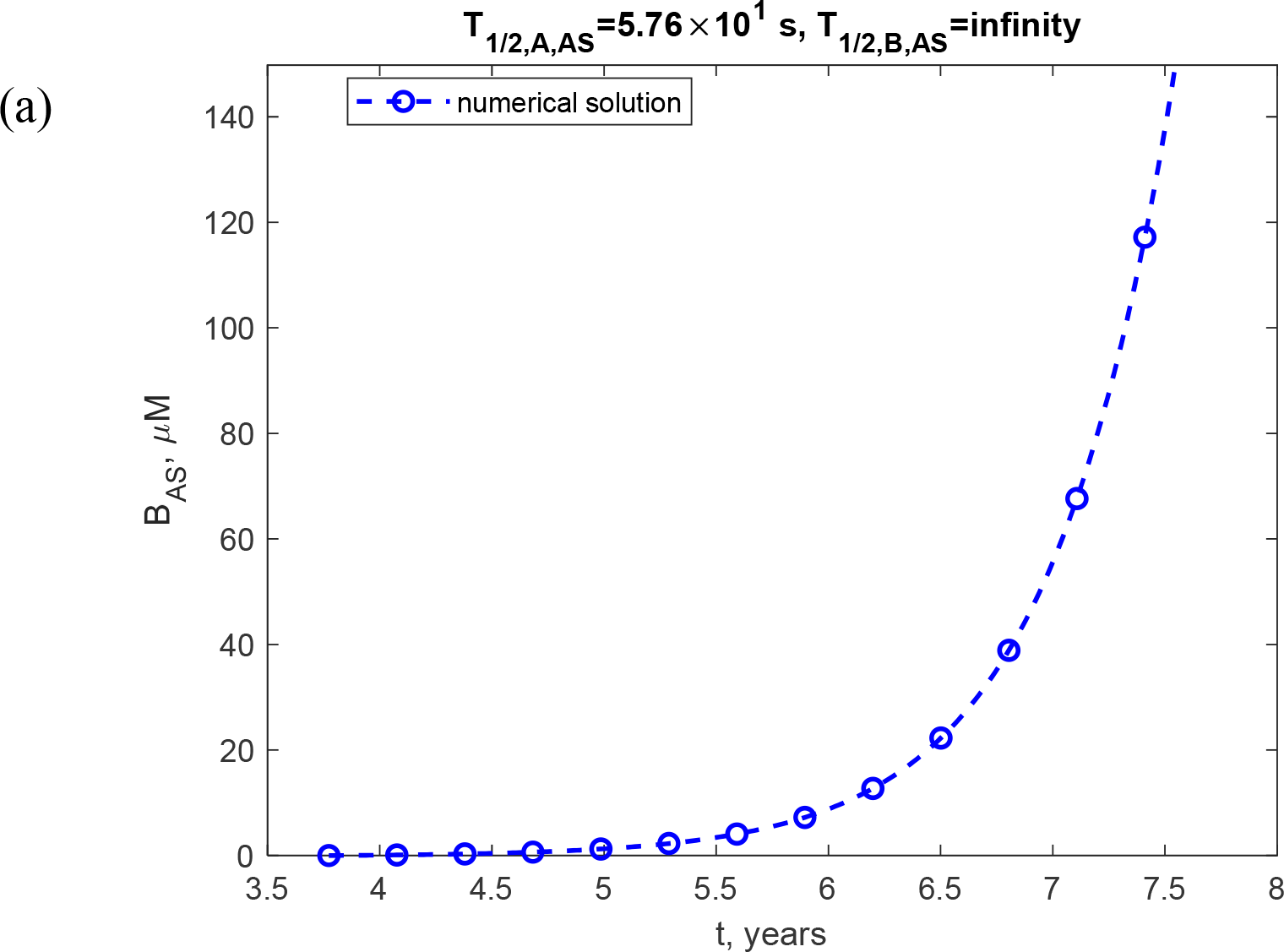
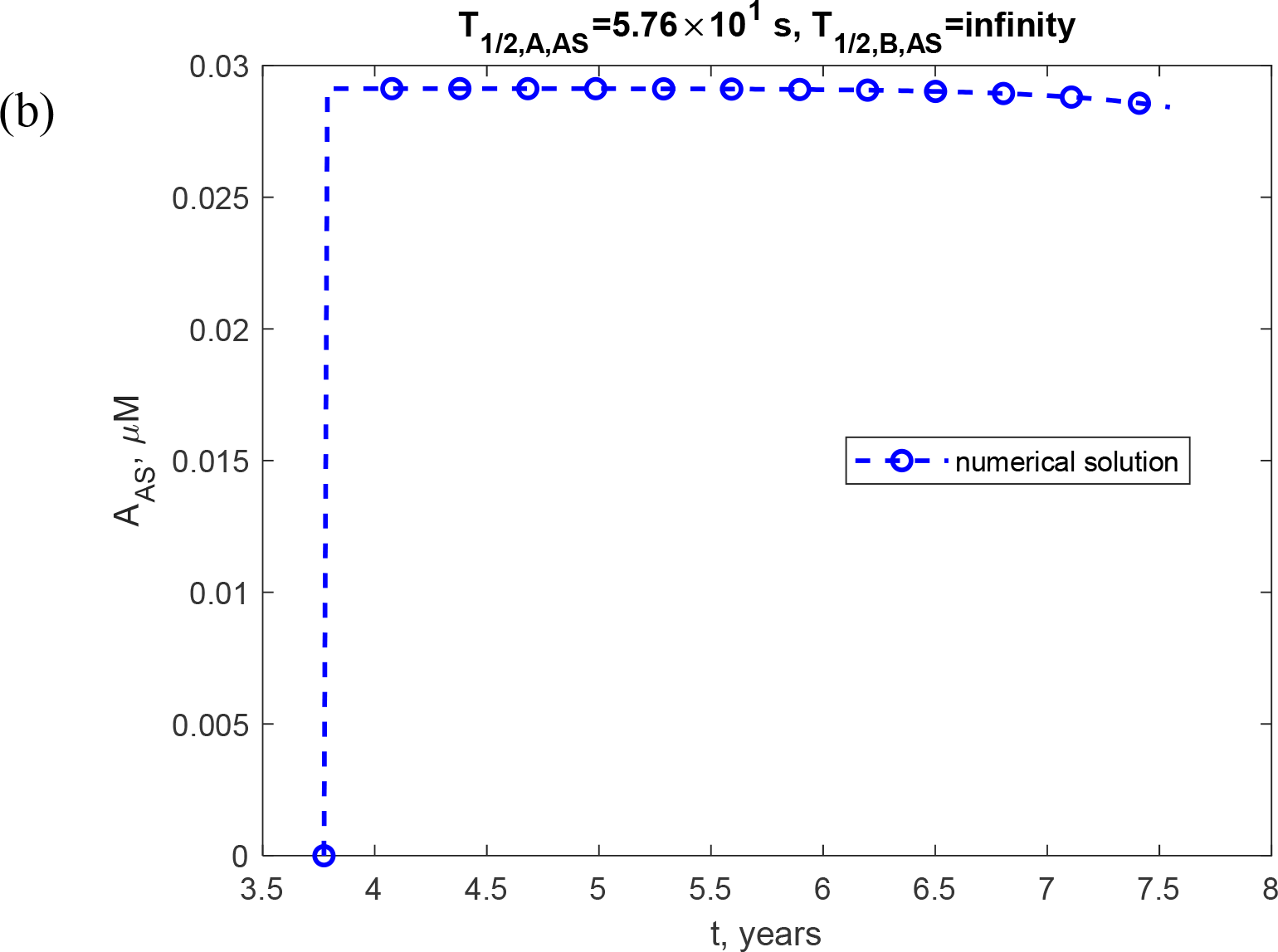
(a) Molar concentration of α-syn fibrils, [*B*_*AS*_] (a) and α-syn monomers, [*A*_*AS*_] (b) as a function of time (in years). This scenario, depicted in the figure, assumes that α-syn monomers have a finite half-life, *T*_1/ 2, *A, FR*_ = 5.76 × 10^1^ s, while α-syn fibrils have an infinite half-life. *q*_*FR*_ =1.57×10^−28^ mol s^-1^ and *q*_*AS*_ =1.47×10^−21^ mol s^-1^.

**Fig. S8.**
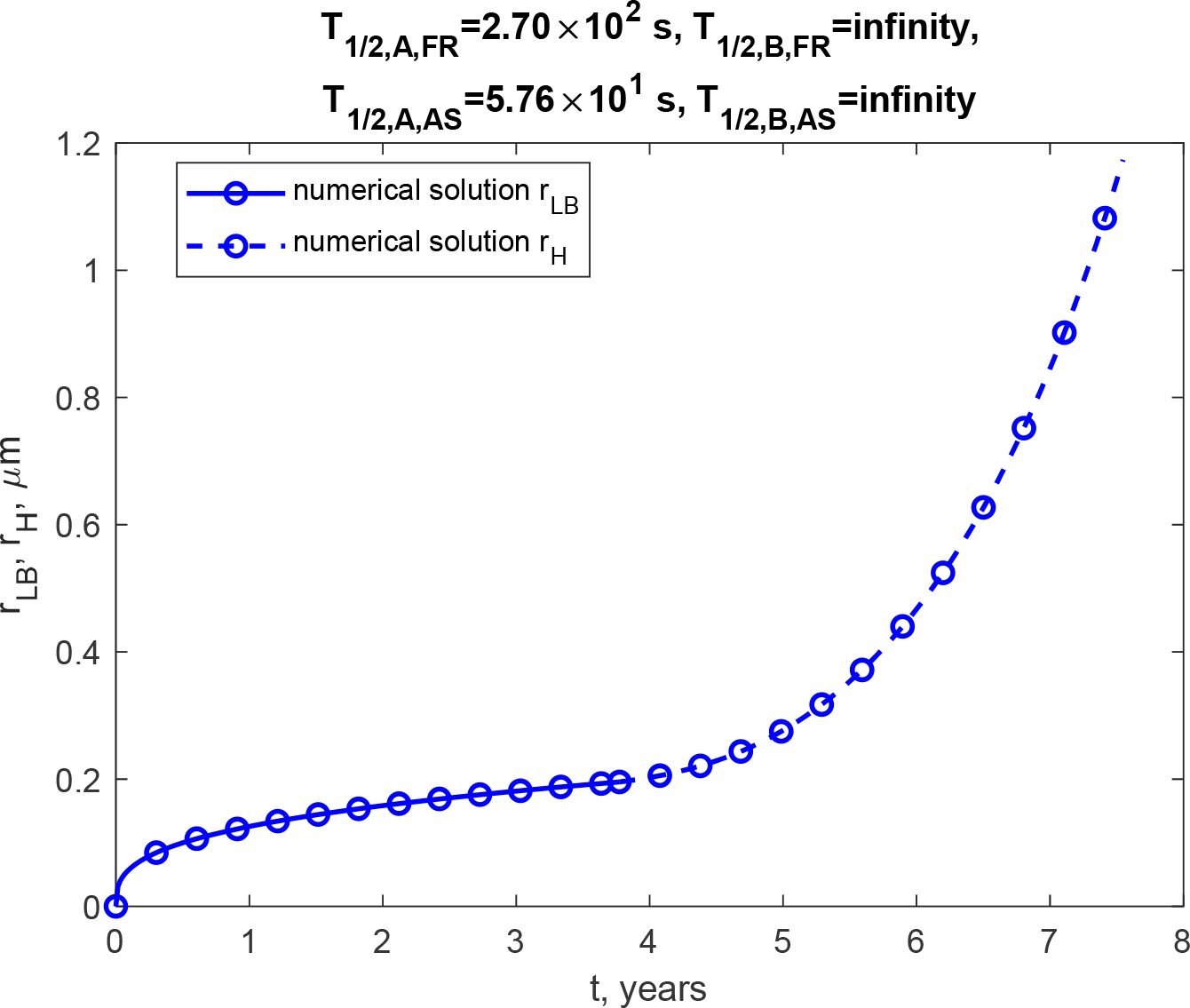
Radius of the growing core of the LB, *r*_*LB*_ and the growing halo, *r*_*H*_ over time (in years). This scenario, depicted in the figure, assumes that membrane fragments and α-syn monomers have finite half-life, *T*_1/ 2, *A, FR*_ = 2.70 × 10^2^ s and *T*_1/ 2, *A, AS*_ = 5.76 ×10^1^ s, while fragment aggregates and α-syn fibrils have infinite half-lives. *q*_*FR*_ =1.57×10^−28^ mol s^-1^ and *q*_*AS*_ =1.47×10^−21^ mol s^-1^.

## Notes

### Competing Interest Statement

The authors have declared no competing interest.

